# Efficient transgenesis and homology-directed gene targeting in monolayers of primary human small intestinal and colonic epithelial stem cells

**DOI:** 10.1101/2021.09.08.459297

**Authors:** Keith A. Breau, Meryem T. Ok, Ismael Gomez-Martinez, Joseph Burclaff, Nathan P. Kohn, Scott T. Magness

## Abstract

**Background & Aims:** 2D monolayers of primary intestinal and colonic epithelial cells represent next-generation *in vitro* models of the gut. Efficient transgenesis and gene-editing in human intestinal stem cells (hISCs) would significantly improve utility of these models by enabling generation of reporter and loss/gain-of-function hISCs, but no published methods exist for transfecting 2D hISC monolayers. Electroporation has proven effective in other difficult-to-transfect cells; thus we applied this method to hISCs.

**Methods:** Twenty-four electroporation parameters were tested, and the optimal condition for efficiency and viability was validated on hISCs from six anatomical regions along the small intestine and colon. PiggyBac™ transposase and Cas9 ribonucleoprotein (RNP) complexes were used for stable genomic integration of reporter genes. High-throughput methods for clone isolation, expansion, and screening were developed. An hISC OLFM4-emGFP reporter was generated and validated by qPCR, organoid assays, and hISC compartmentalization on a planar crypt-microarray (PCM) device.

**Results:** Maximum electroporation efficiency was 79.9% with a mean survival of 65%. Transfection of 10^5^ hISCs produced ∼142 (0.14%) stable transposase-mediated clones. Transfection of *OLFM4*-targetting RNPs yielded ∼35% editing and 99/220 (45%) of antibiotic-resistant colonies analyzed expressed emGFP. OLFM4-emGFP hISCs applied to PCMs remained emGFP+ and proliferative in high-Wnt3a/R-spondin3/Noggin zones yet differentiated to emGFP-/KRT20+ cells outside engineered crypt zones. OLFM4-emGFP levels correlated with endogenous *OLFM4*. Olfm4-emGFP^high^ cells were *LGR5*^high^/*KRT20*^low^, and demonstrated high organoid-forming potential.

**Conclusions:** Electroporation of hISCs is highly efficient for stable transgenesis and transgenic lines can be generated in 3-4 weeks. Workflows mirror conventional culture methods, facilitating rapid integration into established tissue-culture operations. *OLFM4*^high^ is a robust hISC marker with functional properties in culture.

## Introduction

In humans, the lower digestive track is divided into two complementary organs, the small intestine and colon, which carry out different functions to maintain health. Digested food enters the small intestine from the stomach and travels through the lumen, a tubular space that is surrounded by a one cell-thick layer of epithelial cells, which must create a barrier to harmful luminal contents while simultaneously serving as the primary mediator of food and orally-administered drug uptake^1^. Luminal contents then pass to the colonic epithelium, which is the primary mediator of water and electrolyte uptake^2^. The small intestinal and colonic epithelium exist in a highly toxic environment created from substances in food, drink, and oral medications, as well as the commensal and pathogenic bacteria that reside in the lumen. The epithelial cells that are damaged by the luminal contents are rapidly and continually replaced to maintain the broad range of physiological functions and preserve a heathy digestive system.

Physiological turnover of the epithelial tissue occurs about every 5-7 days and is driven by a pool of actively dividing intestinal stem cells (ISCs) located in the base of microanatomical ‘crypt’ regions^3,4^. In the small intestine, ISCs give rise to a larger pool of transit amplifying progenitor cells that migrate out of the crypt onto finger-like projections called villi, terminally differentiating into absorptive cells and minority populations of various secretory cell types^3^. ISCs in the colon are also localized to the crypt base and in general follow a similar migration and differentiation trajectory as in the small intestine, but naturally devoid of villi, the differentiated colonic lineages terminally migrate to a flat epithelial surface^5^. Importantly, disruption of the epithelial barrier or normal ISC renewal in the intestine can lead to a number of human health conditions such as inflammatory bowel disease (IBD) and cancer^6,7,8^. The genetic and cellular mechanisms underlying physiological epithelial renewal is an active area of research and developing better methods and technologies to understand these mechanisms has strong potential to answer basic scientific questions and address health conditions that impair the integrity or function of the gut lining.

The primary means to study the human small intestinal and colonic epithelium has historically been to use fixed tissue, immortalized colon cancer cell lines, or animal models, which in many cases are limited in physiological relevance. While primary human epithelial cells can attach to tissue culture plastic, normal hISCs cannot self-renew in this context, and differentiated epithelial cells are unable to create long-lived confluent cultures due to their inability to proliferate and short life span^9^. Long-term culture of normal primary human epithelium from the small intestine and colon has enormous potential to produce more physiologically relevant models. In the past decade, long-term culture of human small intestinal and colonic epithelium was accomplished through hISC-driven organoid cultures^10,11^. Organoids are small spherical epithelial tissue structures that grow in 3-dimensional hydrogels with media formulations that promote ISC self-renewal by mimicking ISC niche signaling^12^. ISC differentiation can occur stochastically or be induced using native protein factors or small molecules^12^. While organoids address many of the historical short-comings of human gut epithelial culture, their use and throughput for many applications is technically limited by very small size, lack of simultaneous access to the apical and basolateral aspects, and challenges in imaging and quantification through multiple Z-planes of the 3D hydrogels.

The concept of using specialized media and extracellular matrices to recreate an ‘*in vitro* hISC niche’, which enabled 3D organoid cultures, has subsequently been applied to create 2D monolayers of small intestinal and colonic epithelium^9^. To establish monolayers, hISCs are isolated from small patient biopsies or tissue resections and expanded on thick collagen hydrogels with media containing Wnt3a, Noggin, Rspondin-3, EGF and other supportive small molecular factors that support hISC expansion and repress differentiation^9^. 2D hISC culture methods are highly analogous to conventional methods used for immortalized cell lines and thus offer a high level of familiarity and utility. Notably, after expansion as 2D monolayers, hISCs have been directly applied to a number of culture platforms and differentiated to study various aspects of intestinal biology. These platforms include commercially-available permeable Transwell™ inserts, which allow easy differential access to luminal and basolateral aspects of cultured tissues and measurement of barrier function^13,14^, microfabricated devices that re-create the 3D microanatomical features of crypts and villi using biocompatible extracellular matrix scaffolds^15,16^, and ‘2D crypt’ microphysiological systems that zonate hISCs into proliferative and differentiated compartments in a 2D plane^17^.

Given the broad range of emerging uses for colonic and small intestinal hISCs grown as 2D monolayers, the ability to genetically modify hISCs cultured in this manner would significantly improve their utility for disease modeling by facilitating the production of reporter gene lines, gain- and loss-of-function lines, and other sophisticated genetic models. Previous work in 3D colonic organoids compared the efficiency of lentiviral, liposomal, and electroporation transfection methods and demonstrated that electroporation consistently generated the highest percentage of transfected cells, with approximately 30% transfection efficiency and 0.035% PiggyBac™ integration rate^18^. While these methods were successful in achieving gene editing in primary ISCs, they have so far been limited in scope to organoids^18,19,20,21^. Thus, there is a need for efficient transfection methodologies for both small intestinal and colonic hISCs grown as 2D monolayers.

Given that electroporation has generated transfection efficiencies in excess of 75% in other difficult-to-transfect cell lines^22^, and was shown to be the most effective method of transfection in hISC organoids^18,21^, we hypothesized that it could be optimized to efficiently transfect hISCs grown as monolayers. Here we test a wide range of electroporation parameters to define optimal conditions, measure the broad applicability of these parameters on monolayers grown from six regions along the entire length of the human small intestine and colon, assess efficiencies of transgenesis by random integration and site-specific CRISPR/Cas9 gene editing, and develop high-throughput methods for clonal isolation and expansion of transgenic hISCs. We then apply this new method to create an emGFP reporter gene line to detect hISCs *in vitro* and use these hISC-reporter cells to create zonated tissue constructs on a 2D microphysiological model of crypts.

## Results

### A single high voltage pulse efficiently transfects hISCs with high viability

Electroporation is frequently used to transfect primary cells and can result in high transfection efficiency, but this often comes at the expense of cell viability. High cell viability combined with efficient transfection is highly dependent on cell type, buffer system, and the electrical field and pulse conditions^22^; thus, a number of parameters were tested to identify conditions that supported the highest transfection efficiency in hISCs with high viability. hISC cultures were established from crypts isolated from three anatomical regions of the small intestine (duodenum, jejunum, and ileum) from a single organ transplant donor to allow for direct comparison of transfection parameters without donor-to-donor variability. Prior to electroporation, crypts were seeded onto expansion culture plates that contain defined hydrogels and media formulations that support the expansion of hISCs and repress differentiation^9,23^. Low passage (p<12) hISCs were dissociated to single cells, and the Neon™ electroporation system was used to transfect aliquots of 2×10^5^ cells with a constitutively expressing EGFP (Enhanced Green Fluorescent Protein) plasmid providing a visual readout of electroporation efficiency. Twenty-four electroporation conditions that varied in voltage, number of pulses, and pulse duration were tested (Fig. 1A and Supplemental Table 1). Following electroporation, hISCs were recovered on collagen-coated 24-well plates for 48 hours, which provided sufficient time for cell recovery and expression of EGFP.

**Figure 1.**
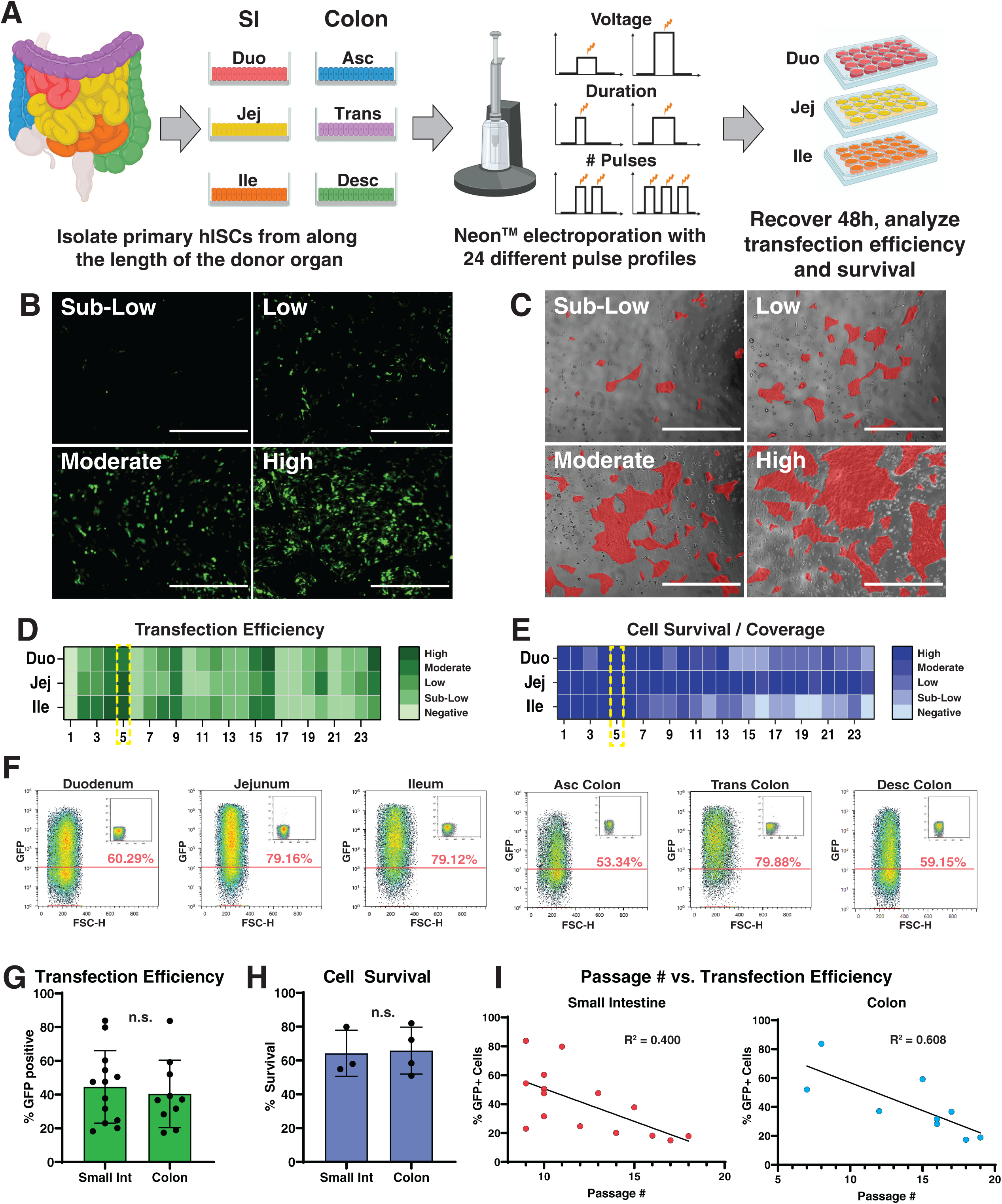
A single high-voltage pulse efficiently transfects hISCs. **(A)** Workflow for hISC sourcing and electroporation optimization. Colors represent different isolated regions of the intestinal tract. Neon™ electroporation system was used to test voltage, pulse duration, and number of pulses; **(B)** Representative images of cell densities used to bin transfection efficiency. Cultures were imaged and scored based on the percentage of GFP+ cells estimated by microscopy: negative (no visible GFP, not pictured), sub-low (<10% of cells expressing GFP), low (10-30%), moderate (30-50%), and high (>50%); Scale bars 2000 μm; **(C)** Representative images of cell densities used to bin cell survival and coverage. For easy visualization, cell patches are overlaid in red. Cell survival was estimated as percentage coverage compared to an untransfected control: negative (no surviving cells, not pictured), sub-low (<10% coverage compared to negative control), low (10-30%), moderate (30-50%), and high (>50%); Scale bars 2000 μm; **(D)** Heatmap summary of transfection efficiencies from 24 tested electroporation parameters with optimal parameters highlighted. Darker color indicates higher transfection efficiency; **(E)** Heatmap summary of cell survival/coverage from 24 tested electroporation parameters with optimal parameters highlighted. Darker color indicates higher cell survival; **(F)** Flow cytometry density plots demonstrating highest transfection efficiency observed for each of six intestinal segments using parameter #5 (1700 V, 20 ms, 1 pulse). Insets show matched untransfected controls for each transfection; **(G)** Summary of flow cytometry-measured transfection efficiencies using parameter #5 (1700 V, 20 ms, 1 pulse); *p = 0*.*96*, n>10 biological replicates per region; **(H)** Summary of hemocytometer-measured cell survival after transfection as a percentage of cell numbers in matched untransfected controls; *p = 0*.*88*, n=3 biological replicates per region; **(I)** Transfection efficiency (from Fig. 1G), plotted as a function of passage number with line of best fit.

A semi-quantitative approach was used to rapidly assess and filter transfection efficiencies from the 72 independent electroporations. Transfected hISCs in 24-well plates were visually evaluated for EGFP expression (i.e. transfection efficiency) and binned into five categories based on estimated percent of EGFP expressing cells: negative (no visible GFP), sublow (<10%), low (10-30%), moderate (30-50%), and high (>50%) (Fig. 1B,D). Cell survival was scored in a similar manner by visually comparing cell coverage in a 24-well plate to an untransfected negative control 48 hours after transfection (Fig. 1C,E). Analyses of all conditions yielded a single electroporation condition (1700 V, 20 ms, 1 pulse) that consistently yielded high transfection efficiency and viability in cells from all from three regions of the small intestine (Fig. 1D).

To quantify and extend these findings, this optimized electroporation condition was next evaluated for its efficacy for transfecting hISCs from six anatomical regions of the small intestine (duodenum, jejunum, and ileum) and colon (ascending, transverse, and descending regions). Flow cytometric analysis of transfected hISCs demonstrated the optimized electroporation parameter was capable of high transfection efficiencies that ranged from ∼59% to ∼80% (Fig. 1F) between different regions. Twenty-three independent electroporations over different passage numbers indicated the mean transfection efficiency to be 40.6% ± 20.94 with a cell survival of 65.2% ± 12.6 and no significant differences between small intestine and colon hISCs (Fig. 1G,H). When transfection efficiencies were compared with hISC passage number, there was a negative correlation (R^2^ = 0.400 for small intestine and 0.608 for colon), suggesting that transfection efficiencies are highest with early passage cells (Fig. 1I). Analysis of only low-passage (p<12) transfections indicated a mean transfection efficiency of 56.7% ± 20.89 in these cells.

### New methods for transgenic colony isolation from 2D collagen hydrogels are fast and high-throughput

Isolation and expansion of clonal cell populations following transfection is a critical step in developing transgenic hISC lines. Because transgenic hISC colonies are grown on the thick collagen hydrogels required for hISC self-renewal, new methods had to be developed for efficient isolation and expansion of individual intact colonies. To develop these methods, antibiotic resistant colonies were generated by electroporating a PiggyBac™ transposase plasmid into hISCs. Upon expression, PiggyBac transposase binds to a pair of ITRs flanking a DNA cassette of interest, and randomly integrates that DNA into the genome. Generation of antibiotic-resistant colonies was thus achieved by co-electroporating a plasmid expressing PiggyBac transposase with a second donor plasmid containing a pair of inverted terminal repeats (ITRs) that flanked a puromycin resistance cassette. Following electroporation, hISCs were allowed to recover for 3 days followed by puromycin treatment for 6 days (Fig. 2A,B).

**Figure 2.**
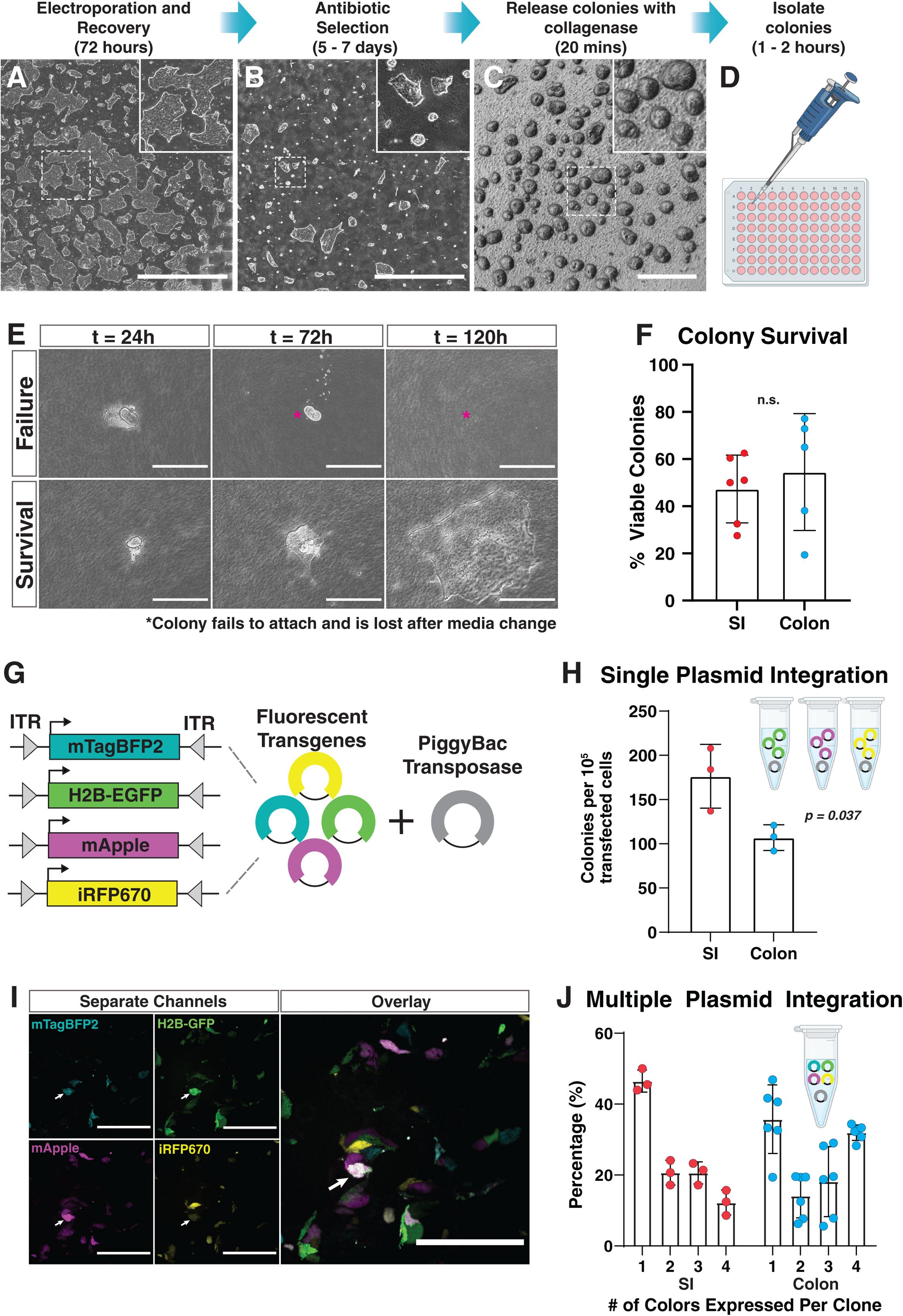
Colony isolation from 2D collagen monolayers is fast and high throughput. **(A-D)** Overview of transfection and colony isolation workflow showing representative images from each stage of the process; Scale bars (A-B) 4000 μm, (C) 400 μm; **(E)** Representative time course of colony attachment and growth after isolation. Asterisk indicates original colony position of non-adherent colony; Scale bars 200 μm; **(F)** Quantification of colony survival following isolation; *p = 0*.*56*, n>4 biological replicates per organ; **(G)** Schematic of plasmids used for quantifying PiggyBac™ transposase integration rate. Each plasmid includes a constitutively-expressed fluorophore flanked by transposase-binding inverted tandem repeats (ITRs); **(H)** Quantification of colony numbers after antibiotic selection for PiggyBac™ integration, normalized by FACS-measured transfection efficiency; *p = 0*.037, n=3 parallel transfections of different plasmids for each organ; **(I)** Representative images of hISCs transfected with the four fluorescent reporter plasmids (Fig. 2G). Arrow indicates a representative colony that has integrated all four reporter plasmids; Scale bars 200 μm; **(J)** Quantification of multiple integration events observed in stably transfected colonies as represented in Fig. 2I. Each data point represents the percentage of a clone type in a different field of view from a single experiment.

To isolate individual clones, a 20-minute incubation in 500 U/mL collagenase IV was used to release the contacts between the hISC colony and the underlying collagen structure (Fig. 2C), releasing intact colonies into the media. Liberated colonies were rinsed in PBS, then transferred into a 10 cm dish for clonal isolation. Individual colonies were collected by pipetting under a bright-field microscope and placed into wells of a collagen-coated 96-well plate containing media for hISC expansion (Fig. 2D). Using this approach, 96 colonies could be isolated in ∼45 minutes. 96-well plates were fed every other day and monitored by microscopy for attachment and growth (Fig. 2E). Four days after clonal isolation, 50.6% ± 19.1 of the isolated hISC colonies demonstrated attachment and increased cell numbers by visual inspection. (Fig. 2F).

### Transfection methods efficiently generate stable single- and multiple-transgene integration events using the PiggyBac transposase system

Stable integration of transgenes is a standard approach for generating reporter gene and gain-of-function cell lines. While single integration events may be sufficient for some applications, multiple integration events are often desired to increase payload expression or express multiple genes in the same cell. To evaluate the efficiency of both single- and multiple-integration events using our method, we generated four different PiggyBac donor vectors, each containing a different constitutively expressed fluorescent protein (mTagBFP2, H2B-EGFP, mApple, and iRFP670) cDNA cassette (Fig. 2G), selected for their very low spectral overlap. First, three of these plasmids were individually mixed with PiggyBac transposase plasmid and transfected into separate aliquots of ∼10^6^ well-dissociated hISCs. Stable single-plasmid integration was quantified after antibiotic selection by counting the number of well-isolated fluorescent colonies. To test the throughput of colony generation, these six transfections were performed in parallel using both jejunal and descending colon monolayers, generating an estimated total of 1,700 antibiotic-resistant colonies with stable fluorescence. To account for variability in transfection efficiency between experiments, the transfection efficiency for each different plasmid was normalized to the efficiency of a control transfection using a constitutively-expressing EGFP plasmid. Following normalization, the rate of integration was determined to be 176 ± 36 colonies (0.18%) per 10^5^ transfected cells in jejunum and 107 ± 15 (0.11%) colonies per 10^5^ transfected cells in descending colon (Fig. 2H), approximately 3-to-4-fold higher than previous reports of PiggyBac integration efficiency in organoids^18^.

To quantify the integration of multiple plasmids in the same cell, equal amounts of each plasmid were mixed and electroporated into hISCs from the jejunum and descending colon, and drug resistant colonies were evaluated under fluorescence microscopy to identify one or more of the four fluorescent protein colors, indicating the number of different plasmids that were integrated (Fig. 2I). For jejunal hISCs, single plasmid integration events occurred in ∼45% of colonies, integration of 2 or 3 different plasmids occurred in ∼20% of clones, and integration of all 4 plasmids occurred in ∼10% of colonies (Fig. 2J). In descending colon hISCs, single plasmid integration events occurred in ∼35% of colonies, integration of 2 or 3 different plasmids occurred in ∼15-20% of clones, and integration of all 4 plasmids occurred in ∼30% of colonies (Fig. 2J). These findings indicated multiple constructs can be efficiently integrated into the genome of the same cell.

### Optimized electroporation parameters are efficient for transfecting CRIPSR/Cas9 RNP complexes

An alternative approach to random transgene integration is CRISPR-based gene editing, which produces highly specific integration into targeted loci and generation of insertion or deletion mutants for gain- and loss-of-function studies^24,25,26^. Transfecting complexes of ribonucleoproteins (RNPs) instead of Cas9-expressing plasmids and guide RNAs independently is an attractive method for delivering CRISPR reagents due to the reduced potential for off-target cleavage or integration^27^. As a candidate locus to test RNP targeting efficiency in hISCs, *OLFM4* was chosen because it is robustly expressed in hISCs of the small intestine, as demonstrated by *in situ* hybridization, immunostaining, and single-cell RNA sequencing in humans^28,29,30^. A guide RNA (gRNA) targeting the 3’ UTR of *OLFM4* was designed using CCTop (which identifies and ranks gRNAs according to predicted off-target activity)^31^ and selected based on minimal risk for off-target cleavage (Supplemental Table 2). Assembled Cas9/gRNA RNPs were electroporated into jejunal and descending colon hISCs. Following recovery, DNA was extracted in bulk from transfected cells and the *OLFM4* locus was sequenced (Supplemental Table 3). TIDE analysis, which detects and quantifies insertion/deletion (indel) mutations by sequencing^32^, demonstrated 34.8% ± 13.4 indel-formation efficiency at this locus with no significant difference between small intestine and colon hISCs (Fig. 3A). These findings show that RNPs are an efficient alternative to plasmid-dependent Cas9 expression for gene editing in 2D monolayers.

**Figure 3.**
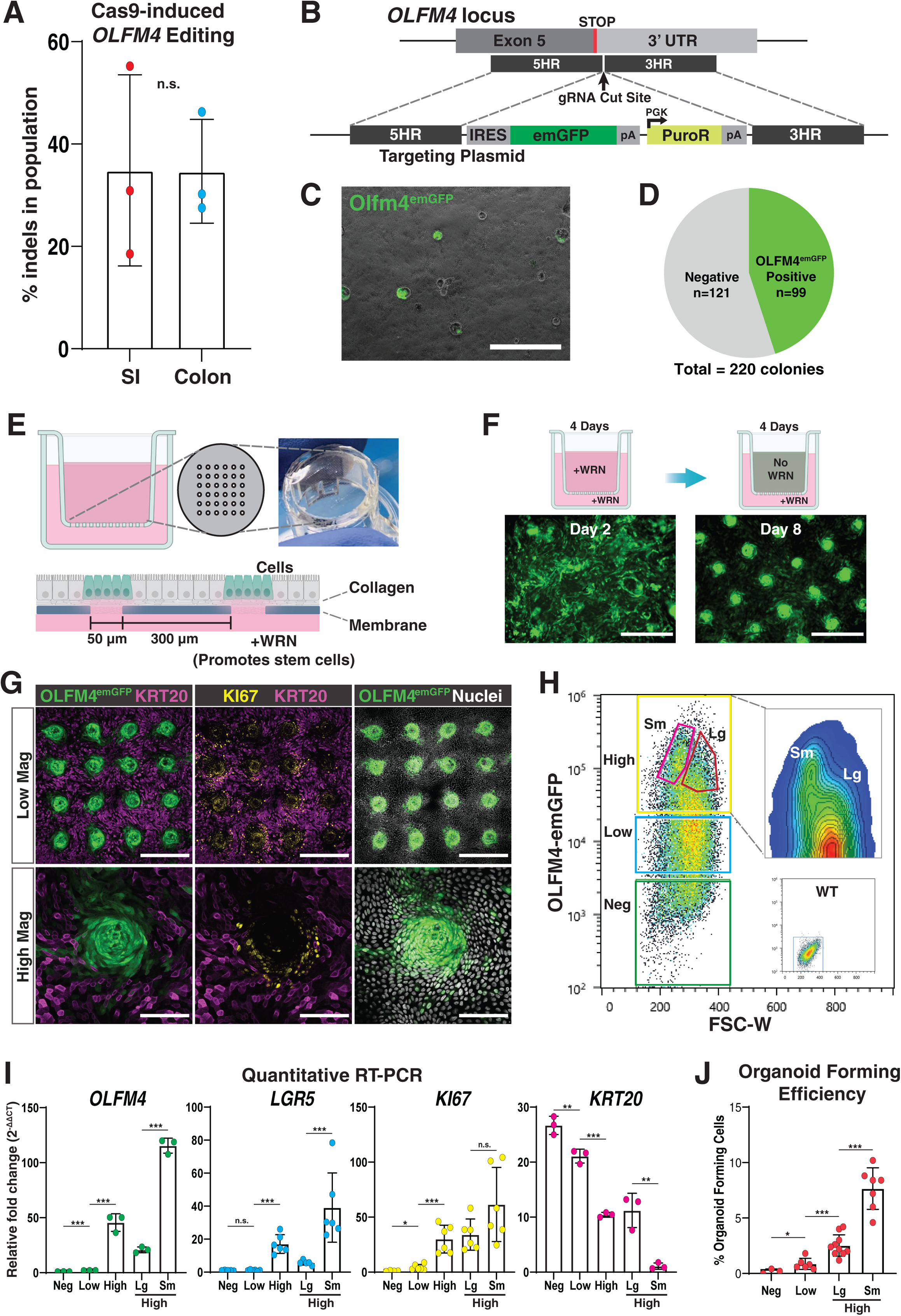
An OLFM4^emGFP^ reporter generated by Cas9-mediated HDR reports proliferative stem cell populations *in vitro*. **(A)** Percentage of Cas9-induced indels at the *OLFM4* gRNA cleavage site, measured by TIDE analysis of bulk DNA collected 5 days after transfection of Cas9/gRNA complexes; *p = 0*.99, n=3 biological replicates per organ; **(B)** Schematic of the *OLFM4* insertion site and plasmid used for targeting, showing IRES-emGFP insertion in the OLFM4 3’ UTR along with a constitutively-expressed puromycin resistance gene; **(C)** Representative image of OLFM4^emGFP^ colonies following antibiotic selection; Scale bar 1000 μm; **(D)** Quantification of fluorescent OFLM4^emGFP^ colonies from Fig. 3C compared to total colony numbers; **(E)** Planar crypt-microarray (PCM) device used for hISC zonation. An array of 50 μm microholes in an impermeable membrane restricts cell access to the WRN-containing basal media reservoir, promoting stem cell maintenance only over the holes; **(F)** Overview of growth and zonation protocol for hISCs grown on PCMs. Cells are grown 4 days with WRN in both apical and basal compartments followed by 4 days with WRN in only the basal compartment; Scale bars 200 μm; (**G)** Confocal microscopy of zonated OLFM4^emGFP^ hISCs grown on PCMs, fixed and stained with antibodies for KRT20 and KI67. Nuclei counterstained with Bisbenzimide A; Scale Bars: Low Mag 400 μm, High Mag 100 μm; **(H)** FACS density plot of OLFM4^emGFP^ cells isolated from a zonated PCM, overlaid with gates used for sorting. Insets show WT negative control and a contour plot of the ‘High’ gate; **(I)** Relative gene expression measured by qRT-PCR on FACS-sorted cells obtained from each gate shown in Fig. 3H. Data normalized to ‘Neg’ for *OFLM4, LGR5*, and *KI67*, and to ‘High^sm^’ for *KRT20*; *\* p < 0*.*05, * * p < 0*.*01, * * * p < 0*.*001;* **(J)** Quantification of organoid formation by FACS-isolated single cells obtained from zonated PCMs; *\* p<0*.*05, * * * p<0*.001, each data point represents the percentage of cells that formed organoids following seeding of 500 single cells into a separate Matrigel™ patty.

These RNP complexes were then used to generate an OLFM4-emGFP reporter line in duodenal hISCs by co-electroporation with a donor plasmid containing an IRES-emGFP cassette flanked by ∼1 kb of WT sequence to facilitate integration through homology-directed repair (HDR, Fig 3B). By integrating into the 3’-UTR of the *OLFM4* locus, the IRES-emGFP cassette reports the presence of *OLFM4* mRNA without disrupting the *OLFM4* coding sequence (Fig. 3B). RNP complexes were electroporated into ∼10^6^ hISCs, and visual inspection by bright field microscopy estimated ∼367 antibiotic resistant clones were present in the well. Visual inspection by fluorescent microscopy also confirmed emGFP expression in ∼45% of colonies (Fig. 3C,D). Forty-eight clonal colonies were isolated and expanded. Five emGFP+ clones were evaluated by PCR, confirming that all exhibited proper integration of the IRES-emGFP cassette in the 3’ UTR of *OLFM4* (Supplemental Fig. 1). One clone (clone D7, Supplemental Fig. 1) was chosen for functional validation. These data demonstrate efficient HDR-mediated gene integration and robust reporter expression from the *OLFM4* locus, highlighting its utility as an hISC reporter gene.

### Olfm4-emGFP hISCs generated by Cas9-mediated transgenesis display compartmentalized GFP expression on a planar crypt-microarray (PCM) device

Since multiple colonies showed ubiquitous OLFM4-emGFP expression and PCR-validated insertion into the 3’-UTR of the *OLFM4* locus, we next sought to validate that the transgene accurately reflected expression of endogenous *OLFM4*. To do this, a recently developed PCM or ‘2D crypt array’ system was employed. The PCM is a microfabricated culture surface that replaces the fully-permeable membrane of a Transwell™ insert with an impermeable membrane possessing permeable holes arranged in a grid-like pattern across its surface^17^. The dimensions of the permeable regions mimic the scale and distance between crypts observed *in vivo*. Tissue zonation, defined as the compartmentalization of proliferative and differentiated zones, is forced by application of hISC maintenance media to the basal reservoir and differentiation media to the apical reservoir (Fig. 3E). hISC functional properties are only maintained over permeable holes in the culture surface providing access to the hISC maintenance factors (basal reservoir), while impermeable zones lacking access to the hISC maintenance factors differentiate (Fig. 3E).

The OLFM4-emGFP hISC reporter line was applied to a PCM and the cells were first allowed to expand as hISCs by applying media with high levels of hISC maintenance factors Wnt3a, R-Spondin3, and Noggin (WRN) on both the apical and basal side of the device (Fig. 3F). After the hISCs reached confluence, the hISC media in the apical reservoir was replaced with differentiation media which lacks hISC factors (Fig. 3F). After 4 days the tissue compartmentalized into OLFM4-emGFP positive and negative zones (Fig. 3F,G). OLFM4-emGFP was strongly expressed over the permeable zones but not in the intervening regions (Fig. 3F,G) suggesting functional ISC and differentiated cell zonation. Immunostaining for the general proliferation marker (KI67) showed proliferative cells (consistent with hISC/progenitors)^33^ localized primarily over the permeable crypt zones (Fig. 3G). By contrast, Cytokeratin 20 (KRT20), a pan-marker of differentiated intestinal epithelial cells^34,35^, showed localization primarily in the intervening impermeable regions (Fig. 3G). Consistent with proper ISC zonation, confocal microscopy demonstrated robust co-localization of OLFM4-emGFP and Ki67 expression over the permeable crypt zones and visibly low or no transgene expression in the intervening zones (Fig. 3G). These data demonstrate that the OLFM4-emGFP reporter cells retain characteristics of hISCs in that they respond to self-renewal signaling and can differentiate into post-mitotic lineages. Additionally, these findings suggest that the OLFM4-emGFP reporter can be used to distinguish between functional ISCs and differentiated cells in culture.

### Olfm4-EGFP cells mimic endogenous *OLFM4* expression and allow for isolation of *LGR5*^high^/*KRT20*^low^ cells with high organoid-forming potential

Flow cytometric analysis of cells zonated on the PCM demonstrated a range of OLFM4-emGFP expression levels that were classified as negative, low (bottom 50% of EGFP+ cells), and high (top 50% of GFP+ cells). Gates for FACS (Fluorescence-Activated Cell Sorting) were defined based on these three expression categories (Fig. 3H). Interestingly, the forward-scatter (FSC-W) parameter indicated a separate density of cells within the OLFM4-emGFP^high^ population that suggested a population of cells with a smaller diameter than the main distribution of cells (High^Sm^, Fig. 3H). Additional FACS gates were included to compare the apparently smaller population of OLFM4-emGFP^high^ cells with the apparently larger OLFM4-emGFP^high^ cells based on the forward-scatter parameter (High^Lg^, Fig. 3H). To verify that the OLFM4-emGFP signal was consistent with endogenous *OLFM4* expression, cells from each gate were FACS-isolated and qPCR was performed on cDNA generated from each population (Fig. 3I). The extent and magnitude of OLFM4-emGFP transgene expression was consistent with endogenous *OLFM4* expression (Fig. 3I). The expression of hallmark ISC and proliferation markers, *LGR5* and *KI67*, tracked with increasing *OLFM4* and the OLFM4-emGFP transgene (Fig. 3I). Conversely, the pan-differentiated cell marker gene, *KRT20*, negatively correlated with the ISC and proliferation marker genes (Fig. 3I). Interestingly, the smaller OLFM4-emGFP^high^ population (High^Sm^) demonstrated significantly higher levels of hISC marker genes compared to the larger OLFM4-emGFP^high^ population (High^Lg^) (Fig 3I). These results show that cells zonated on PCMs demonstrate a range of *OLFM4* mRNA expression, and suggest functional differences associated with both OLFM4-emGFP expression and cell size.

The capacity for single cells to generate multi-cellular organoids has been previously used to functionally demonstrate stemness in ISCs^12^. To investigate the relationship between OLFM4-emGFP expression and proliferative capacity, single cells from each FACS-gated population were sorted into Matrigel™ patties and assayed for their ability to form organoids. Following seeding, individual cell diameters were measured, confirming the smaller diameter of the High^Sm^ population suggested by forward-scatter values (Supplemental Fig. 2). Organoids that developed from single cells were quantified after seven days of growth. The results show that <1% of cells from the GFP-negative and GFP-low gates formed organoids, while 2.52% and 7.66% of the cells from the High^Lg^ and High^Sm^ gates, respectively, formed organoids (Fig. 3J). These data demonstrate a strong positive correlation between OLFM4-emGFP expression and organoid forming capacity, and moreover demonstrate that the smaller OLFM4-EGFP^high^ cells have heightened ISC capacity in culture (Fig. 3J).

## Conclusions

The ability to rapidly expand small samples of human intestinal epithelium as 2D monolayers holds enormous potential for studying intestinal disease. Consequently, the ability to generate genetic models from these cultures would significantly extend their utility, enabling diverse models ranging from reporter lines for screening drug interactions in the small intestine to genetic knockout models of IBD in the colon. In this paper we present new methods for the generation and isolation of transgenic small intestinal and colonic hISCs grown as 2D monolayers. We optimize electroporation parameters and demonstrate that these parameters are capable of robustly transfecting hISCs isolated from six representative regions of the intestinal tract with a high rate of cell survival. We show that these parameters efficiently transfect both DNA, in the form of PiggyBac™ plasmids, and protein, in the form of Cas9 RNP complexes, and demonstrate their utility by generating a novel and robust OLFM4-emGFP reporter line for hISCs.

Our data demonstrates a broad range of transfection efficiencies ranging from 18% to 79% even using optimized parameters. While the exact mechanisms mediating this variability are not well understood, a number of associated factors have been identified that may contribute. Cell passage numbers has been shown in other reports to impact transfection efficiencies^36,37^. Our findings are consistent with this as there was a correlation between decreased transfection efficiency and increasing cell passage number. This highlights an important experimental design parameter to be considered. It has also been shown that proliferative state and cell density can affect transfection efficiency^38^. These parameters were not explored in our study as the transfection efficiencies were more than sufficient for most applications.

Another unexplored possibility is instability of media components used for hISC culture. As described in many reports, hISC culture is commonly carried out using incompletely-defined medium. ISCs rely on a number of heat- and time-labile factors including Wnt3a, R-spondins, Noggin, and EGF^10^. Wnt3a, a potent hISC maintenance factor, must be palmitoylated for high biological activity^39^, thus Wnt3a, as well as Noggin and R-spondins are routinely supplied to hISC cultures using media conditioned from a tripartite mammalian cell line with stable transgenes that express murine Wnt3a, Noggin, and R-spondin3 and also efficiently palmitoylates transgenic Wnt3a (L-WRN cells)^40,41^. While this medium has been shown to produce consistent amounts of Wnt activity, it is still unknown how prolonged culture of these cells impacts the concentrations and biological activities of growth factors and other metabolites. To further highlight uncontrolled variables that could impact electroporation efficiencies, L-WRN cells (and consequently hISCs), are grown in fetal bovine serum (FBS)^9,40^. FBS is known to have high lot-to-lot and vendor-to-vendor variability in growth factor content^42^, and it has been previously suggested that this variability in growth factors may affect hISC growth^23^. Thus, further optimization and quality control of well-defined medias could be performed to improve the consistency of transfection.

A common challenge in random-integration transgenesis and targeted gene editing is isolating populations of cells that are truly clonal, that is, come from a single gene-edited cell. While confirmation of this is ultimately dependent on post-hoc clone validation, the likelihood of obtaining clonal populations is significantly improved by having well-isolated colonies following antibiotic selection. Good experimental design to achieve well-isolated clones is highly dependent on transfection efficiency, payload integration efficiency, and the number of transfected cells. This could be readily observed in PiggyBac-mediated transgenic colonies versus CRISPR/Cas9-HDR mediated transgenesis. The high transfection efficiency and high integration of PiggyBac-mediated transgenesis yielded a large number of antibiotic-resistant clones that merged as they grew as visualized by expression of different fluorescent reporters in small clones that had merged into a single large colony (Fig. 2I). By contrast, the lower efficiency of homology-directed repair (HDR) yielded far less antibiotic-resistant clones that were well-separated (Fig. 3C). Thus, when clonal isolation is necessary, the mode of transgenesis, efficiency of integration, and post-electroporation seeding density should all be considered during experimental design.

While previous publications have presented methods for transfecting hISCs grown as organoids^18,21^, large-scale application of these methods are impeded by significant technical limitations. For example, organoid cultures are grown in a 3D Matrigel™ patty, which has size and volume limitations due to the requirement for growth factors to readily diffuse throughout the gel. This limits the Matrigel volume to ∼25 μL and thus limits the number of organoids that can be expanded in each well. Consequently, organoid-based methods recommend growing upwards of 24 patties to obtain sufficient cell numbers for a single targeted knock-in experiment^18^. By contrast, our data demonstrates that hISCs grown as 2D monolayers can generate hundreds of clones from a single culture well of hISCs, even with a relatively rare event such as HDR-mediated knock-in. Since the personnel time required to passage organoids and 2D hISC monolayers is nearly equivalent in our experience, the methods described here can thus be estimated to produce a 24-fold increase in throughput over existing methods. Furthermore, organoid-based methods require that clones isolated after transfection be individually isolated, dissociated, and embedded in new Matrigel patties, a process which requires a significant time investment. By contrast, isolating colonies from 2D monolayers requires only the amount of time it takes to pipet a colony under the microscope and deposit it into a well, a process that could be readily automated with a large particle sorter.

Another major advantage of this technology is the ability to transfect Cas9 protein complexes directly, which has not been previously demonstrated in hISCs. This is particularly desirable for genome editing as the transient nature of protein transfection has been shown to reduce off-target Cas9 cleavage, and in some cases can significantly improve the level of on-target Cas9 editing^27^. Furthermore, recent improvements in fusion protein design have led to a rapidly expanding toolkit of Cas9-fusion proteins^26^, which could theoretically be transfected using these methods. These include nuclease-null Cas9 fused to: transcriptional activation complexes for activating gene expression^43^, transcriptional repressor complexes for gene silencing^44^, fluorophores for *in vivo* visualization of chromatin domains^45^, base editors for single nucleotide editing^46^, or chromatin remodeling complexes to either re-program post-mitotic cells^47^ or direct the differentiation of stem cells^48^. A number of recent papers have highlighted significant biological differences between mouse and human intestinal epithelium, including the existence of a completely novel cell type in humans, called BEST4 or BCHE cells^30,49^, and major apparent differences in the role of Paneth cells^30^. As detailed study of these differences will likely require *in vitro* hISC culture, the ability to directly transfect these new Cas9 fusion proteins into hISCs has significant potential in discerning the origin and function of these cells in humans.

As another way of studying hISC differentiation, robust reporter lines for human stem cells are highly sought after. Previous reporters for hISCs have focused on *LGR5*^50^, a WNT target gene which has been demonstrated to specifically mark multipotent crypt-base columnar (CBC) stem cells in murine lineage tracing models^33^. However, single cell RNA sequencing (scRNA-seq) of primary human tissue demonstrates that *LGR5* is expressed at low levels in human ISCs and is highly restricted in its expression to a small number of cells^51^, limiting its potential usefulness in isolating non-CBC progenitor populations. Recent studies highlight a gradient of cellular plasticity in early progenitor populations^52^, and this is an active area of study that would benefit from reporters that allow differential isolation of stem and progenitor populations along this gradient.

In contrast to *LGR5, OLFM4* is a direct target of both NOTCH and WNT signaling^53,54^, and has been shown by both single molecule fluorescence in situ hybridization (smFISH) and scRNA-seq to be robustly expressed by human CBCs, with gradually decreasing expression along the length of the crypt^28,30^. By combining an OLFM4-emGFP reporter with PCMs, our data revealed several novel findings. First, the results demonstrate biocompatibility of human ISCs with the PCM device. Second, FACS analysis shows a robust gradient of OLFM4-emGFP expression levels with a direct correlation between high OLFM4-emGFP expression and high KI67 (proliferative capacity) and shows the ability to isolate hISC states with gradually decreasing stem cell function. Third, findings indicate that cells with the highest OLFM4-emGFP expression and hISC organoid forming capacity also have a smaller cell diameter. While the functional consequences of cell sizes are not well understood, small cell size has been correlated in several stem cell types with high stem cell function^55,56^, thus, the combination of OLFM4-emGFP reporter with PCMs is a powerful system that allows both robust real-time visualization of stem cell populations and facilitates isolation of multiple physiologically relevant populations of hISCs.

In the past decade, advances in genome engineering have revolutionized the way in which we study cellular biology. Consequently, the ability to more easily apply these tools to hISCs has significant potential to expediate intestinal research. In this paper, we show that gene editing in hISCs cultured as 2D monolayers is highly efficient, thus laying the groundwork for generating future genetic models of human intestinal epithelium.

## Methods

### hISC Media Preparation

Maintenance media (MM) for hISC culture was prepared as previously described^23^. Briefly, confluent L cells expressing transgenic Wnt3a, Noggin, and Rspondin3 (https://www.atcc.org/products/crl-3276) were cultured in Advanced DMEM/F12 (Gibco 12634010) plus 1X GlutaMAX™ (Gibco 35050061), 1X Pen/Strep (Gibco 15070063), and 20% FBS (VWR Premium 97068-085, GeminiBio Foundation™ 900-108) This media was collected and replaced every 24 hours for 12 days. Collected “conditioned” media (CM) was then pooled, filtered, and frozen into aliquots stored at -80° C. Thawed CM aliquots were mixed 1:1 with basal media (BM) consisting of 2% B-27 (Gibco 12587001), 10 mM Nicotinamide (Sigma-Aldrich N0636), 10 mM HEPES (Corning 25-060-Cl), 1X GlutaMAX, 1X Pen/Strep, 1.25 mM N-Acetylcysteine (Sigma-Aldrich A9165), 50 μg/mL Primocin (Invivogen ant-pm-05), 3 μM SB202190 (Peprotech 1523072), 50 ng/mL mEGF (Peprotech 315-09), 2 nM Gastrin (Sigma-Aldrich G9145), and 10 nM Prostaglandin E2 (Peprotech 3632464). For plating of primary crypts and following passaging, MM was supplemented with 10 μM Y27632 (MM+Y, Selleck Chemical S6390).

### Collagen Plate Preparation

Collagen coated plates for monolayer culture were prepared as previously described^9^. Cold rat tail collagen (Corning 354236) was neutralized with a buffer containing: 1X PBS (Gibco 14190-144), 20 mM HEPES, 0.45% NaHCO_3_ (Gibco 25080-094), and NaOH (23 μL of 1N NaOH per 1 mL of collagen stock solution, EM Science 1310-73-2) to a final collagen concentration of 1 mg/mL. 1 mL cold neutralized collagen solution was added to 6-well tissue culture plates (Genesee 25-105) pre-warmed to 37 °C, allowed to polymerize for 1 hour, then overlaid with 3 mL 1X PBS. Plates were rinsed 3 times with 3 mL 1X PBS (5 minutes of room temperature incubation per rinse) before use.

### Tissue Procurement and Dissection

Intestinal tissue was isolated post-mortem from a 12 y.o. Caucasian male with no known history of intestinal disease and negative for an infectious disease panel performed peri-mortem. Whole intestine was obtained post-mortem from HonorBridge (formally Carolina Donor Services). Following trimming of external adipose tissue, intestine length was measured and the first 20 cm of the small intestine proximal to the pancreatic duct was isolated. A 9 cm^2^ tissue sample was taken from the center of this piece (duodenum). The remaining small intestine was divided in half, and tissue samples were taken from the middle of each half (jejunum and ileum). The colon was divided into three equal length pieces, and tissue samples were taken from the center of each piece (ascending, transverse, descending Colon). Tissue samples were stored in Advanced DMEM/F12 plus 10 μM Y27632 and 200 μg/mL Primocin during dissection, which was immediately followed by crypt isolation.

### Crypt Isolation and hISC Expansion

Following dissection, tissue sections were incubated in cold PBS plus 10 mM N-Acetylcysteine with gentle agitation for 15 minutes. They were then transferred to Isolation buffer (IB) consisting of 5.6 mM Na_2_HPO_4_ (Sigma S7907), 8.0 mM KH_2_PO_4_ (Sigma P5655), 96.2 mM NaCl (Sigma S5886), 1.6 mM KCl (Sigma P5405), 43.4 mM Sucrose (Fisher BP 220-1), 54.9 mM d-sorbitol (Fisher BP439-500), and 100 μM Y27632 and rinsed briefly, then transferred to IB plus 2 mM EDTA (Corning 46-034-Cl) and 0.5 mM DTT (Fisher Scientific BP172-5) and incubated for 30 minutes with gentle agitation. Tubes were shaken vigorously for 2 minutes, after which sections were transferred to new tubes of IB+EDTA+DTT and rocked gently for 10 minutes. This process of vigorous shaking, transfer to a new tube, and gentle rocking was repeated six times. Buffer aliquots were analyzed under widefield microscopy to quantify villi and crypt numbers, and crypt-enriched samples were pooled. Crypt numbers were counted, and seeded on collagen plates in hISC expansion media with Y27632 (MM+Y, described above), 200 μg/mL Primocin, 200 μg/mL Gentamycin (Sigma Aldrich G1914), and 0.5 μg/mL Amphotericin B (Sigma-Aldrich A2942) (plus antibiotics) at 5,000 crypts per well of a 6-well plate. Plates were incubated at 37 °C and media was changed daily with fresh MM plus antibiotics.

### Routine hISC Monolayer Culture and Passaging

Human ISCs were cultured at 37 °C, 5% CO_2_ in 3 mL of MM as described above, replacing the media every two days. Cells were passaged 1:3 every 4-7 days based on cell density, with a desired pre-passage confluency of >60%. 10 μM Y27632 was added to each well at least 1 hr before passaging. To passage, 1 mL of media was removed from each well into a conical tube and the remaining media aspirated. Collagen patties were gently dislodged with a pipet tip and transferred to the tube. 100 μL of 5,000 U/mL collagenase IV (Fisher Scientific LS004189) in HBSS (Gibco 14175-095) was added to each tube and the collagen patties were gently broken up by pipetting with a 5 mL serological pipet. This solution was incubated at 37 °C for 20 minutes, mixing gently every 5 minutes, then pelleted at 800 x g for 3 minutes in a clinical centrifuge. Supernatant was then aspirated and the cell pellet resuspended in 5 mL pre-warmed 1X PBS + 10 μM Y27632. PBS suspensions were incubated at 37 °C for 5 minutes, then pelleted at 800 x g for 3 minutes. PBS was aspirated and cell pellets were then resuspended in 200 μL TrypLE Express (Gibco 12605-010) and incubated at 37 °C for 5 minutes. TrypLE solutions were pipetted ∼15 times with a P1000 pipet tip to dissociate cells, then neutralized with 2 mL MM and pelleted again at 800 x g for 3 minutes. Supernatant was aspirated and the cells were resuspended in 9 mL MM+Y per dissociated well and plated onto three new pre-rinsed collagen-coated wells (3 mL/well).

### *OLFM4* Targeting Plasmid Generation and gRNA Selection

Donor-specific genomic DNA was extracted by dissociating cultured monolayers as described above, then resuspending the final cell pellet in proteinase K solution (10 μM Tris-HCl pH 8.0, 10 μM EDTA, 10 μM NaCl, 0.5% Sarkosyl, 1 mg/mL proteinase K) and incubating at 55 °C overnight. DNA was purified by phenol:chloroform extraction followed by isopropanol precipitation, a 70% ethanol wash, and resuspension in TE Buffer (10 mM Tris-Cl, 1 mM EDTA, pH 8.0). *OLFM4* homology regions were amplified with CloneAmp™ HiFi PCR Premix (Takara 639298) using primers KB39+KB40 and KB69+KB38 (Supplemental Table 3), which included 20 bp of overlapping homology to plasmid backbone elements, and purified on a silica minicolumn. IRES-emGFP and vector backbone were isolated from a pre-existing plasmid by restriction digestion, and assembled with PCR-amplified homology arms using In-Fusion® HD Cloning Kit (Takara 638920). Final products were verified by enzymatic digestion and sequencing. The *OLFM4* gRNA was designed using CCTop^31^ and was selected based on location in the proximal 3’ UTR and low probability of off-target cleavage (Supplemental Table 2).

### Preparation of Transfection Reagents

Super PiggyBac™ Transposase Expression Vector was purchased from System Biosciences (PB210PA-1) and added at a ratio of 5 ng/μL of transfection mix. All other plasmids for transfection were prepared from bacterial stocks, incubating 100 mL LB cultures with appropriate antibiotics overnight with vigorous agitation (225 rpm). Plasmid DNA was purified using QIAGEN HiSpeed® Maxi kit (QIAGEN 12662), eluting in 1 mL TE Buffer. Plasmids were then precipitated with isopropanol, rinsed with 70% ethanol, and resuspended at 1-2 μg/μL in TE Buffer. pMax-GFP plasmid (http://www.addgene.org/vector-database/3525/) was added at a ratio of 100 ng/μL of transfection mix. All other plasmids were added at 50 ng/μL. For gRNA and Cas9 transfections, crRNA and tracrRNA oligos were purchased from IDT (Alt-R® CRISPR-Cas9 system) and resuspended at 100 pmol/μL in nuclease-free dH2O (Corning 46-000-Cl). 2 μL (200 pmol) each of gRNA and tracRNA were combined with 2 μL of 5X annealing buffer (ThermoFisher 100061876) and 4 μL nuclease-free dH2O, and annealed in a thermocycler (95 °C – 5 min, 95 °C to 78 °C @ -2 °C/s, 78 °C – 10 min, 78 °C to 25 °C @ -0.1 °C/s, 25 °C – 5 min), then immediately transferred to ice. 3 μL (60 pmol) of this mixture was added to 2 μL of TrueCut™ Cas9 v2 (∼60 pmol, ThermoFisher A36498), incubated at room temperature for 15 minutes to promote formation of Cas9:gRNA complexes, then stored on ice until use.

### Single-Cell Dissociation for Transfection

To improve cell dissociation for transfection, several changes were made to the routine passaging protocol described above. The length of the 37 °C PBS incubation was extended to 10 minutes, with gentle mixing after the first 5 minutes. Next, 20 μL of 5,000 U/mL collagenase IV was added to the 200 μL TrypLE used for dissociation. Following the 5-minute incubation in TrypLE plus collagenase, cells were first dissociated by pipetting ∼15 times with a P1000 pipet tip as above, then further dissociated by gently pipetting with a 28G insulin syringe (BD Biosciences 329424) 5-7 times. Dissociated TrypLE solutions were neutralized, centrifuged, and aspirated normally, but were then resuspended in 1 mL 1X room temperature PBS. 10 μL of this cell suspension was removed and mixed 1:1 with trypan blue and cell numbers were quantified with a hemocytometer. PBS solutions were again pelleted at 800 x g for 3 minutes, the supernatant aspirated, and the cell pellet resuspended in cold Neon™ Buffer R (ThermoFisher MPK10096) at a concentration of 5,000 - 20,000 cells/μL.

### Electroporation and Antibiotic Selection

For electroporation, cells were dissociated to singlets as described above. For testing of electroporation conditions, cells were resuspended at a concentration of 20,000 cells/μL in Neon™ Buffer R, and pMax-GFP was added at a concentration of 100 ng/μL buffer. Following electroporation with a 10 μL Neon tip (ThermoFisher MPK1096), cells were seeded onto a single well of a collagen-coated 24-well plate with MM+Y and allowed to recover overnight at 37 °C. Media was replaced after 24 hours with MM, and cells were analyzed for transfection and survival after 72 hours. After optimization, all other transfections were performed using the Neon Electroporator 100 μL tips (ThermoFisher MPK10096) with preset #5 (1700 V, 1 pulse, 20 ms), electroporating 6,000 – 12,000 cells/μL and plating onto a single collagen-coated well of a 6-well plate. Media was changed after 24 hours with MM, then again with MM plus puromycin (2 μg/mL, Sigma P8833) after 72 hours and every other day thereafter until no more cell death was observed (5-7 days after addition of puromycin).

### Colony Isolation and Expansion

To isolate transgenic colonies following antibiotic selection, 300 μL of 5,000 U/mL collagenase IV was added to each well and incubated at 37 °C for 20 minutes. Supernatant and dislodged colonies were collected and pelleted at 600 x g. Media was aspirated and the cells were gently resuspended in 10 mL PBS + 10 μM Y27632 (PBS+Y) to rinse. Cells were re-pelleted, and suspended in a second aliquot of 15 mL PBS+Y and added to a 10 cm tissue culture dish. This dish was transferred to an inverted light microscope, and colonies were manually pipetted into a collagen-coated 96-well culture plate containing 100 μL MM+Y per well. Following isolation, colony media was changed every other day with MM+Y and monitored for attachment and growth. Upon reaching confluency or after 2 weeks, colonies were serially passaged to collagen-coated 48-well, 12-well, and 6-well culture plates.

### PCM Fabrication

The PCMs were fabricated as previously described^17^. Briefly, an array of 20 × 20 microholes (50 µm in diameter) were micropatterned onto a thin, impermeable 1002F-10 photoresist film using a photomask (Front Range Photomask) and UV photolithography^57^. The 1002F-10 formulation and spin coating parameters (1500 rpm, 30 s) were chosen to achieve a shallow microhole depth of 10 µm. The micropatterned film was then mounted onto a bottomless 12-well Transwell™ insert using 3M double-sided medical tape and coated with 200 µL of 1 mg/mL neutralized rat tail collagen I (Corning 354236) as previously described. Collagen-coated PCMs were incubated in a humidified 37 °C chamber for 1 hour to promote formation of a uniform gel. The collagen gel was then dried overnight in a 40 °C oven then stored at room temperature. Before seeding cells, PCM Transwells were sterilized with ethanol, then rehydrated with 50 µg/mL rat tail collagen I in PBS overnight at 37 °C.

### Growing hISCs on PCM Devices and Immunofluorescent Staining

Confluent 6-well monolayers of OLFM4-emGFP cells were dissociated as previously described, passaging one well of a 6-well plate to 3 PCM devices in MM+Y, adding 500 μL of medium with dissociated cells to the upper compartment and 1.5 mL of medium (without cells) to the lower compartment. Media was changed after 48 hours with fresh MM+Y. On the fourth day after seeding, medium in the upper compartment was replaced with differentiation media (DM), comprised of Advanced DMEM/F12, 1X GlutaMAX, 1.25 mM N-Acetylcysteine, 10 mM HEPES, 50 ng/mL mEGF, 50 μg/mL Primocin, and 500 μM A83-01 (Sigma-Aldrich SML0788), and medium in the lower compartment was replaced with MM (no Y). Media was changed daily for four consecutive days in this manner (DM/MM). On the eighth day after seeding, cells were either isolated for FACS (see below) or rinsed with PBS and fixed with 4% PFA (FisherScientific AC416780250) for 15 minutes. Following fixation, cells were rinsed with PBS, permeabilized with 0.5% Triton X-100 (FisherScientific AC215682500) in PBS for 20 minutes at room temperature, then rinsed twice with PBS plus 3% BSA (FisherScientific BP1600-1, diluted in PBS). Blocking was performed for 30 minutes at room temperature in 3% BSA. After blocking, cells were incubated with KRT20 primary antibody (1:200 dilution, Cell Signaling D9Z1Z) in 3% BSA for 48 hours at 4 °C, rinsed with 3% BSA, then incubated with a donkey anti-rabbit Cy3 secondary antibody (Jackson ImmunoResearch 711-166-152) and APC-conjugated KI67 antibody (Invitrogen 17-5698-83) for one hour at room temperature. Cells were rinsed twice with 3% BSA, and nuclei were stained with Bisbenzimide A (10 μg/mL, SigmaAldrich B1155) in 3% BSA for 5 minutes, then rinsed and stored in 3% BSA for microscopic analysis.

### FACS, qPCR, and Organoid Formation

Following growth and compartmentalization of PCM devices, culture media was aspirated and replaced with 4 mL warm PBS+Y, incubating for 5 minutes at 37 °C. PBS was then aspirated, and 500 μL warm TrypLE was added to both the upper and lower compartments. Devices were incubated for 10 minutes at 37 °C, pipetting vigorously after 5 and 10 minutes. Dislodged cells were then further dissociated with a 28G insulin syringe, and TrypLE was neutralized by adding 4 mL of culture media. Cells were pelleted at 800 x g for 3 minutes, resuspended in 1 mL of PBS then re-pelleted. Cells were then resuspended in MM+Y and sorted by FACS using a Sony SH800ZF cell sorter (SN 1500005). FSC-A, BSC-A, FSC-H, and BSC-H parameters were used to filter out small debris and cell doublets, enriching for live single cells, and FSC-H and GFP parameters (GFP sensor gain 32.5%) were used to sort populations expressing different levels of OLFM4-emGFP into cold MM+Y. For qPCR, RNA was isolated using the RNAqueous™ Micro Kit (Invitrogen AM1931). cDNA amplification and qPCR were performed with SsoAdvanced™ Universal Probe Supermix (Bio-Rad 1725281), using TaqMan probes Hs00197437_m1 (*OLFM4*), Hs00969442_m1 (*LGR5*), Hs04260396_g1(*MKI67*), Hs00300643_m1 (KRT20), and Hs03003631_g1 (*18S*). To quantify organoid formation, 5,000 single cells (2,500 cells for negative population) were sorted into separate tubes and resuspended in 10 μL of ice-cold GFR Matrigel™ (Corning 354230) per 500 cells. 10 μL aliquots of cells were dispensed into separate wells of a 96-well culture plate, incubated for 25 minutes at 37 °C to facilitate gel formation, then overlaid with 100 μL MM+Y. Media was changed every other day with fresh MM for 7 days, at which point organoids were fixed with pre-warmed 4% PFA for 15 minutes for analysis.

### Statistics

Statistical analyses were performed using two-tailed *t*-tests. Most analyses utilized the Student’s *t*-test, with the exception of organoid forming efficiency (Figure 3J), which used Welch’s *t*-test due to the expectation of unequal variance between samples. For comparison of qPCR data, statistics were computed by comparing Δ18S values between samples using a Student’s *t*-test.

## Supporting information

Figures (Full Resolution)

## Acknowledgments

First and foremost, we would like to thank the anonymous organ donor, their family, and HonorBridge (formerly Carolina Donor Services), Organ Procurement Organization, Durham, NC, for providing the tissue samples used in this study. We would also like to thank UNC’s Human Pluripotent Cell Core Facility, for generous use of their Neon™ Transfection instrument, UNC’s Advanced Analytics Core for help with flow cytometry, and the UNC Animal Models Core for providing various DNA elements used in plasmid construction. Figure schematics were created with BioRender.com. This work was performed in part at the Chapel Hill Analytical and Nanofabrication Laboratory, CHANL, a member of the North Carolina Research Triangle Nanotechnology Network, RTNN, which is supported by the National Science Foundation, Grant ECCS-2025064, as part of the National Nanotechnology Coordinated Infrastructure, NNCI. This work was funded by NIH grants T32GM133364, R01DK115806, P30DK034987, R43DK125155, R01DK109559, and the Katherine Bullard Charitable Trust. Scott Magness would like to disclose a financial interest in Altis Biosystems Inc., which licenses the hISC monolayer technology used in this study.

---

**Supplemental Table 1.**
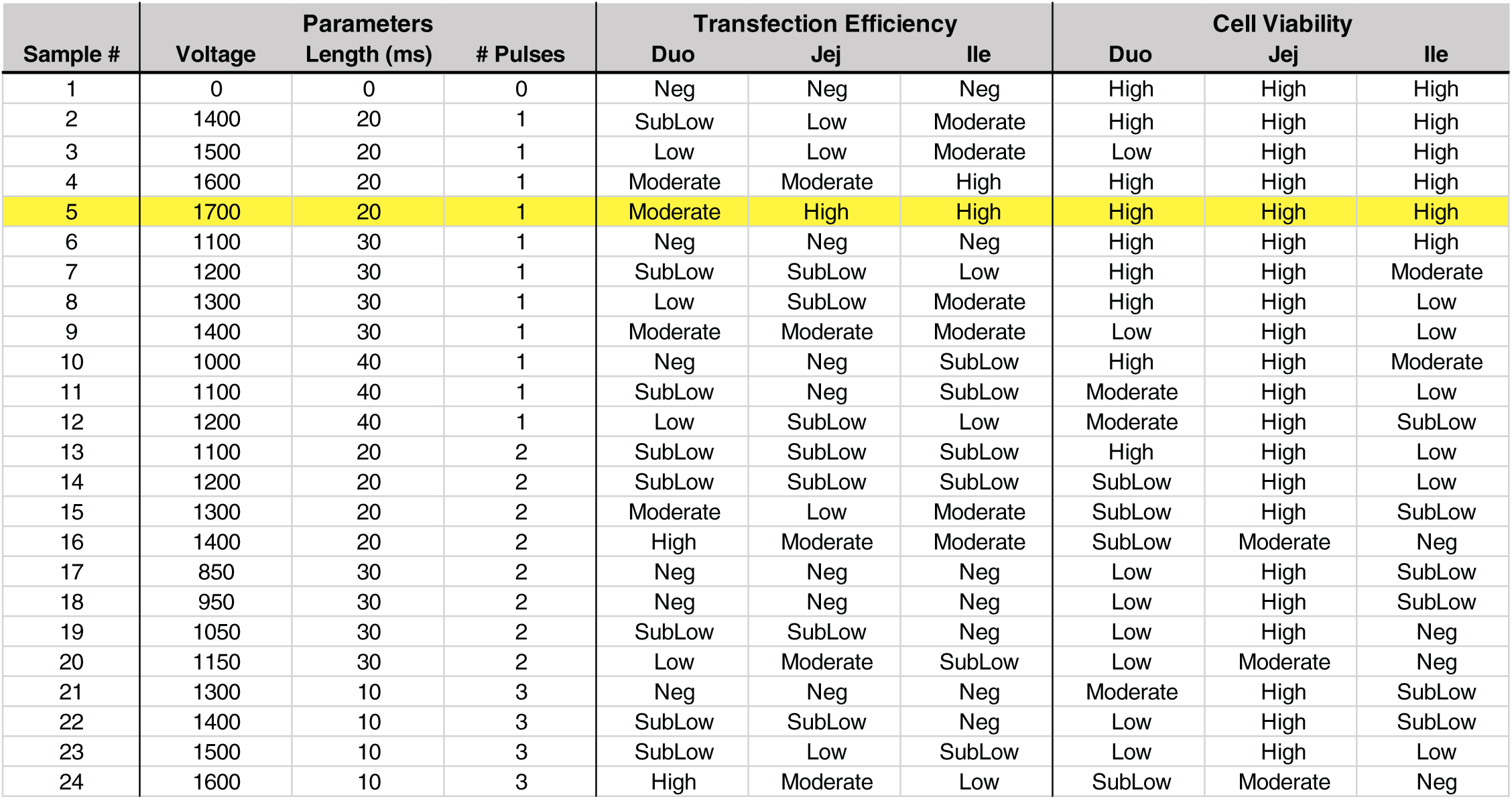
Electrical pulse parameters for optimization. Electrical parameters used to optimize electroporation efficiency. Twenty-four sets of parameters were tested, varying electrical voltage, pulse duration, and number of pulses. Transfection efficiency and cell viability of three biological replicates were estimated and binned into five categories (Neg, SubLow, Low, Moderate, and High). Condition #5 (1700 V, 20 ms, 1 pulse), highlighted, was used for further experiments.

**Supplemental Table 2.**
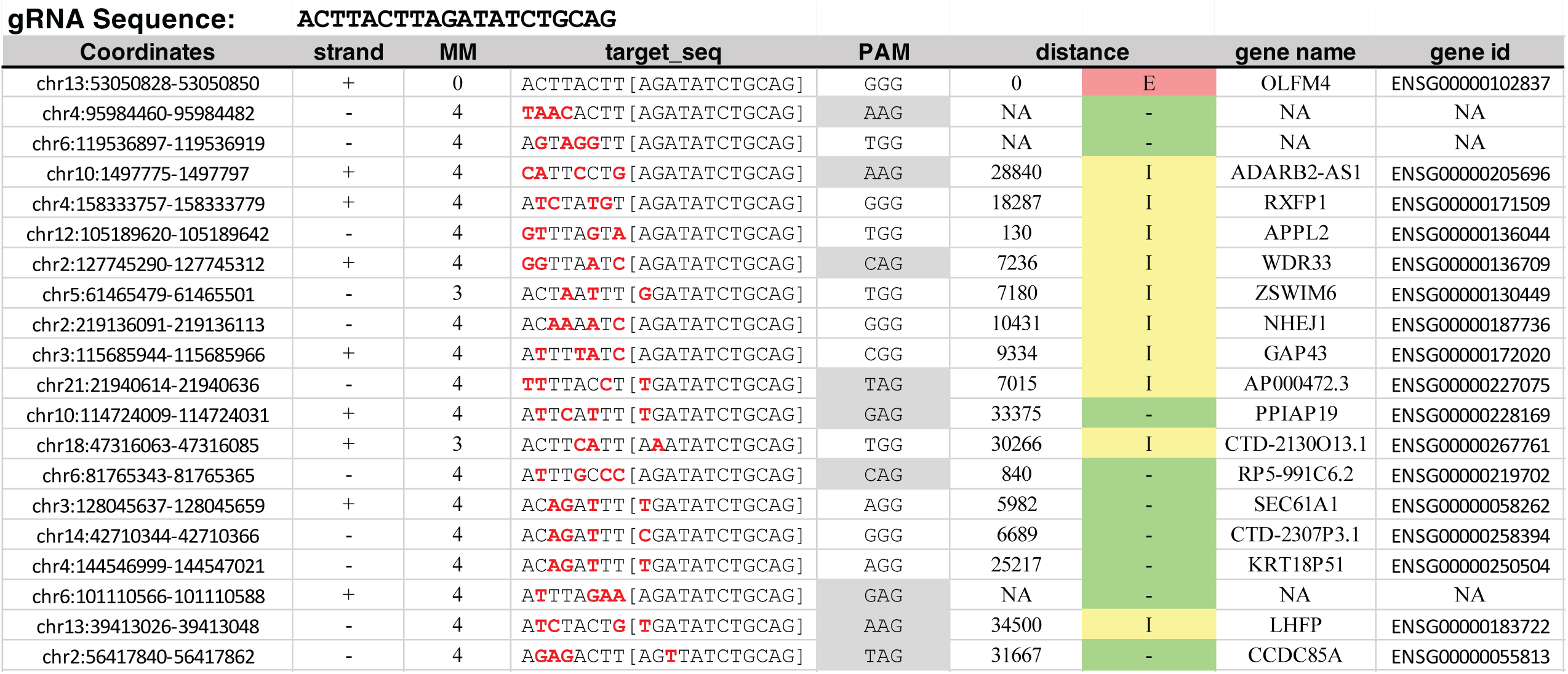
CCTop off-target loci for *OLFM4* gRNA. CCTop output of the selected *OLFM4*-targetting gRNA, showing DNA sequences throughout the genome with homology to the gRNA sequence. First entry is the desired cleavage site in the terminal exon of *OLFM4*. Latter entries are potential off-target cleavage sites, ranked by decreasing predicted risk of cleavage, with mismatched nucleotides shown in red. gRNA was chosen such that all potential off-target sites have at least four mismatches, or three mismatches with at least one mismatch within the 12 bp “seed” region (shown in brackets), proximal to the Cas9 cleavage site.

**Supplemental Table 3.**
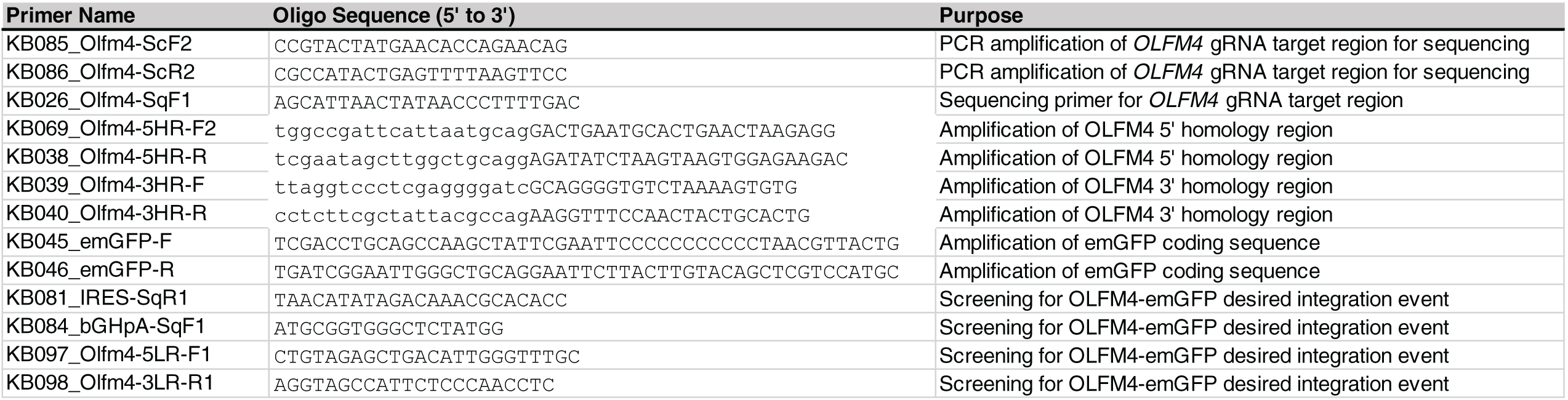
Primers used in this study. Primers used for generating TIDE results (top), generating the *OLFM4* targeting plasmid (middle), and validating integration of the IRES-emGFP sequence (bottom).

**Supplemental Figure 1.**
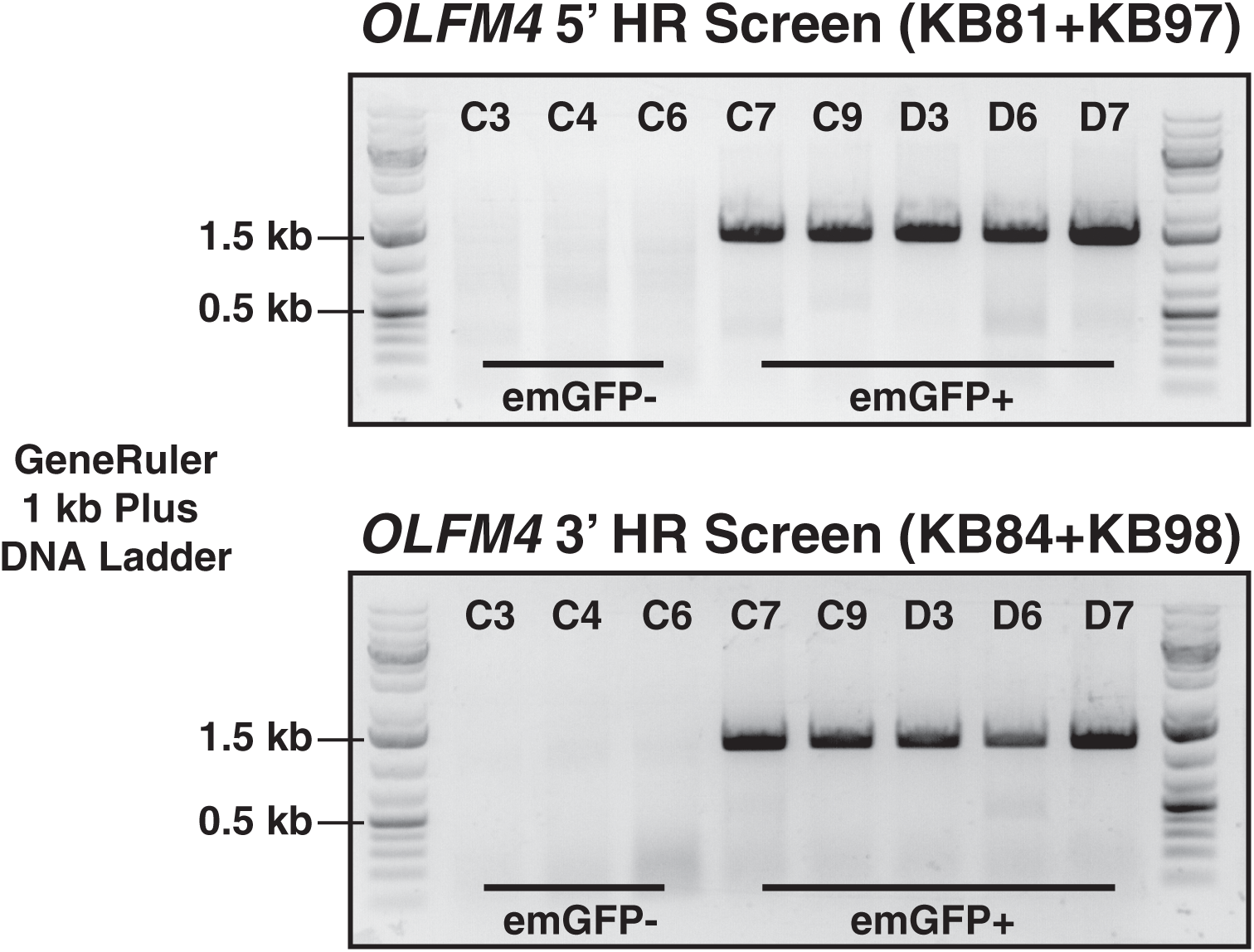
PCR validation of IRES-emGFP integration at the *OLFM4* locus. Three emGFP-negative clones were included for a negative control, along with five emGFP-positive clones. Primers were designed to span the homology arms used for HDR, with one primer in each screen binding to the native *OLFM4* locus outside of the homology regions and the other binding to the IRES-emGFP insert.

**Supplemental Figure 2.**
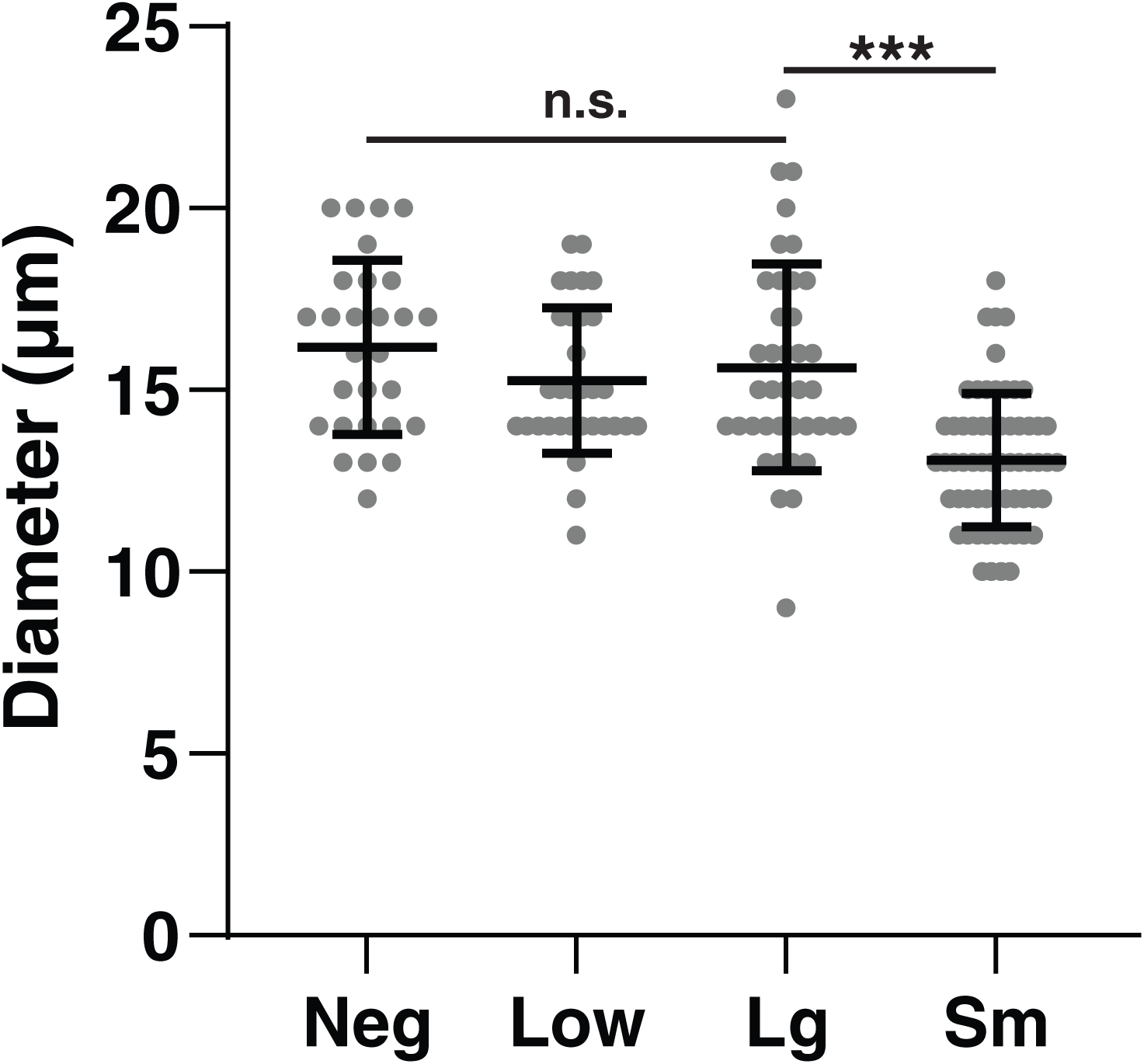
Microscopy-measured cell diameters of FACS-isolated hISCs retrieved from PCM devices and embedded in Matrigel. Each data point represents a different cell. emGFP-High^Sm^ cells display significantly reduced cell size.

## References

1) Peterson, L. W. & Artis, D. Intestinal epithelial cells: regulators of barrier function and immune homeostasis. Nat Rev Immunol 14, 141–153 (2014).

2) Kiela, P. R. & Ghishan, F. K. Physiology of Intestinal Absorption and Secretion. Best practice & research. Clinical gastroenterology 30, 145 (2016).

3) Cheng, H. & Leblond, C. P. Origin, differentiation and renewal of the four main epithelial cell types in the mouse small intestine I. Columnar cell. American Journal of Anatomy 141, 461–479 (1974).

4) Barker, N. Adult intestinal stem cells: critical drivers of epithelial homeostasis and regeneration. Nat Rev Mol Cell Biol 15, 19–33 (2014).

5) Magney, J. E., Erlandsen, S. L., Bjerknes, M. L. & Cheng, H. Scanning electron microscopy of isolated epithelium of the murine gastrointestinal tract: morphology of the basal surface and evidence for paracrinelike cells. Am J Anat 177, 43–53 (1986).

6) Guan, Q. A Comprehensive Review and Update on the Pathogenesis of Inflammatory Bowel Disease. J Immunol Res 2019, 7247238 (2019).

7) Dekker, E., Tanis, P. J., Vleugels, J. L. A., Kasi, P. M. & Wallace, M. B. Colorectal cancer. Lancet 394, 1467–1480 (2019).

8) Cives, M. & Strosberg, J. R. Gastroenteropancreatic Neuroendocrine Tumors. CA Cancer J Clin 68, 471–487 (2018).

9) Wang, Y. et al. Self-renewing Monolayer of Primary Colonic or Rectal Epithelial Cells. Cellular and Molecular Gastroenterology and Hepatology 4, 165-182.e7 (2017).

10) Sato, T. et al. Long-term Expansion of Epithelial Organoids From Human Colon, Adenoma, Adenocarcinoma, and Barrett’s Epithelium. Gastroenterology 141, 1762–1772 (2011).

11) Jung, P. et al. Isolation and in vitro expansion of human colonic stem cells. Nat Med 17, 1225–1227 (2011).

12) Sato, T. et al. Single Lgr5 stem cells build crypt-villus structures in vitro without a mesenchymal niche. Nature 459, 262–265 (2009).

13) Moon, C., VanDussen, K. L., Miyoshi, H. & Stappenbeck, T. S. Development of a primary mouse intestinal epithelial cell monolayer culture system to evaluate factors that modulate IgA transcytosis. Mucosal Immunol 7, 818–828 (2014).

14) VanDussen, K. L. et al. Development of an enhanced human gastrointestinal epithelial culture system to facilitate patient-based assays. Gut 64, 911–920 (2015).

15) Wang, Y. et al. A microengineered collagen scaffold for generating a polarized crypt-villus architecture of human small intestinal epithelium. Biomaterials 128, 44–55 (2017).

16) Wang, Y. et al. Formation of Human Colonic Crypt Array by Application of Chemical Gradients Across a Shaped Epithelial Monolayer. Cellular and Molecular Gastroenterology and Hepatology 5, 113–130 (2018).

17) Kim, R. et al. Formation of arrays of planar, murine, intestinal crypts possessing a stem/proliferative cell compartment and differentiated cell zone. Lab on a Chip 18, 2202–2213 (2018).

18) Fujii, M., Matano, M., Nanki, K. & Sato, T. Efficient genetic engineering of human intestinal organoids using electroporation. Nat Protoc 10, 1474–1485 (2015).

19) Schwank, G. & Clevers, H. CRISPR/Cas9-Mediated Genome Editing of Mouse Small Intestinal Organoids. Methods Mol Biol 1422, 3–11 (2016).

20) Kashfi, H., Jinks, N. & Nateri, A. S. Generating and Utilizing Murine Cas9-Expressing Intestinal Organoids for Large-Scale Knockout Genetic Screening. Methods Mol Biol 2171, 257–269 (2020).

21) Matano, M. et al. Modeling colorectal cancer using CRISPR-Cas9-mediated engineering of human intestinal organoids. Nat Med 21, 256–262 (2015).

22) Jordan, E. T., Collins, M., Terefe, J., Ugozzoli, L. & Rubio, T. Optimizing Electroporation Conditions in Primary and Other Difficult-to-Transfect Cells. J Biomol Tech 19, 328–334 (2008).

23) Hinman, S. S., Wang, Y., Kim, R. & Allbritton, N. L. In vitro generation of self-renewing human intestinal epithelia over planar and shaped collagen hydrogels. Nat Protoc 16, 352–382 (2021).

24) Cong, L. et al. Multiplex genome engineering using CRISPR/Cas systems. Science 339, 819–823 (2013).

25) Mali, P. et al. RNA-guided human genome engineering via Cas9. Science 339, 823–826 (2013).

26) Pickar-Oliver, A. & Gersbach, C. A. The next generation of CRISPR–Cas technologies and applications. Nat Rev Mol Cell Biol 20, 490–507 (2019).

27) Liang, X. et al. Rapid and highly efficient mammalian cell engineering via Cas9 protein transfection. J Biotechnol 208, 44–53 (2015).

28) van der Flier, L. G., Haegebarth, A., Stange, D. E., van de Wetering, M. & Clevers, H. OLFM4 Is a Robust Marker for Stem Cells in Human Intestine and Marks a Subset of Colorectal Cancer Cells. Gastroenterology 137, 15–17 (2009).

29) Gersemann, M. et al. Olfactomedin-4 is a glycoprotein secreted into mucus in active IBD☆. Journal of Crohn’s and Colitis 6, 425–434 (2012).

30) Busslinger, G. A. et al. Human gastrointestinal epithelia of the esophagus, stomach, and duodenum resolved at single-cell resolution. Cell Reports 34, 108819 (2021).

31) Stemmer, M., Thumberger, T., del Sol Keyer, M., Wittbrodt, J. & Mateo, J. L. CCTop: An Intuitive, Flexible and Reliable CRISPR/Cas9 Target Prediction Tool. PLoS One 10, e0124633 (2015).

32) Brinkman, E. K., Chen, T., Amendola, M. & van Steensel, B. Easy quantitative assessment of genome editing by sequence trace decomposition. Nucleic Acids Research 42, e168–e168 (2014).

33) Barker, N. et al. Identification of stem cells in small intestine and colon by marker gene Lgr5. Nature 449, 1003–1007 (2007).

34) Moll, R. et al. Identification of protein IT of the intestinal cytoskeleton as a novel type I cytokeratin with unusual properties and expression patterns. J Cell Biol 111, 567–580 (1990).

35) Merlos-Suárez, A. et al. The Intestinal Stem Cell Signature Identifies Colorectal Cancer Stem Cells and Predicts Disease Relapse. Cell Stem Cell 8, 511–524 (2011).

36) McCoy, A. M., Collins, M. L. & Ugozzoli, L. A. Using the gene pulser MXcell electroporation system to transfect primary cells with high efficiency. J Vis Exp 1662 (2010).

37) de Carvalho, T. G. et al. A simple protocol for transfecting human mesenchymal stem cells. Biotechnol Lett 40, 617–622 (2018).

38) Prasanna, G. L. & Panda, T. Electroporation: basic principles, practical considerations and applications in molecular biology. Bioprocess Engineering 16, 261–264 (1997).

39) Willert, K. et al. Wnt proteins are lipid-modified and can act as stem cell growth factors. Nature 423, 448–452 (2003).

40) Miyoshi, H. & Stappenbeck, T. S. In vitro expansion and genetic modification of gastrointestinal stem cells in spheroid culture. Nat Protoc 8, 2471–2482 (2013).

41) VanDussen, K. L., Sonnek, N. M. & Stappenbeck, T. S. L-WRN conditioned medium for gastrointestinal epithelial stem cell culture shows replicable batch-to-batch activity levels across multiple research teams. Stem Cell Research 37, 101430 (2019).

42) Honn, K. V., Singley, J. A. & Chavin, W. Fetal bovine serum: a multivariate standard. Proc Soc Exp Biol Med 149, 344–347 (1975).

43) Perez-Pinera, P. et al. RNA-guided gene activation by CRISPR-Cas9–based transcription factors. Nat Methods 10, 973–976 (2013).

44) Mandegar, M. A. et al. CRISPR Interference Efficiently Induces Specific and Reversible Gene Silencing in Human iPSCs. Cell Stem Cell 18, 541–553 (2016).

45) Khan, A. O., Simms, V. A., Pike, J. A., Thomas, S. G. & Morgan, N. V. CRISPR-Cas9 Mediated Labelling Allows for Single Molecule Imaging and Resolution. Sci Rep 7, 8450 (2017).

46) Chen, L. et al. Programmable C:G to G:C genome editing with CRISPR-Cas9-directed base excision repair proteins. Nat Commun 12, 1384 (2021).

47) Liu, P., Chen, M., Liu, Y., Qi, L. S. & Ding, S. CRISPR-Based Chromatin Remodeling of the Endogenous Oct4 or Sox2 Locus Enables Reprogramming to Pluripotency. Cell Stem Cell 22, 252-261.e4 (2018).

48) Gong, J., Tang, D., Leong, K.W., CRISPR/dCas9-mediated cell differentiation, Current Opinion in Biomedical Engineering (2018).

49) Parikh, K. et al. Colonic epithelial cell diversity in health and inflammatory bowel disease. Nature 567, 49–55 (2019).

50) Fujii, M. et al. Human Intestinal Organoids Maintain Self-Renewal Capacity and Cellular Diversity in Niche-Inspired Culture Condition. Cell Stem Cell 23, 787-793.e6 (2018).

51) Wang, Y. et al. Single-cell transcriptome analysis reveals differential nutrient absorption functions in human intestine. Journal of Experimental Medicine 217, e20191130 (2020).

52) de Sousa e Melo, F. & de Sauvage, F. J. Cellular Plasticity in Intestinal Homeostasis and Disease. Cell Stem Cell 24, 54–64 (2019).

53) Liu, W. et al. Olfactomedin 4 deletion induces colon adenocarcinoma in ApcMin/+ mice. Oncogene 35, (2016).

54) VanDussen, K. L. et al. Notch signaling modulates proliferation and differentiation of intestinal crypt base columnar stem cells. Development 139, 488–497 (2012).

55) Kucia, M. et al. A population of very small embryonic-like (VSEL) CXCR4(+)SSEA-1(+)Oct-4+ stem cells identified in adult bone marrow. Leukemia 20, 857–869 (2006).

56) Paiva, C. S. D., Pflugfelder, S. C. & Li, D.-Q. Cell Size Correlates with Phenotype and Proliferative Capacity in Human Corneal Epithelial Cells. STEM CELLS 24, 368–375 (2006).

57) Pai, J.-H. et al. Photoresist with Low Fluorescence for Bioanalytical Applications. Anal. Chem. 79, 8774–8780 (2007).

