## Supplementary figures and images for "Efficient transgenesis and homology-directed gene targeting in monolayers of primary human small intestinal and colonic epithelial stem cells"

### Figures (Full Resolution)

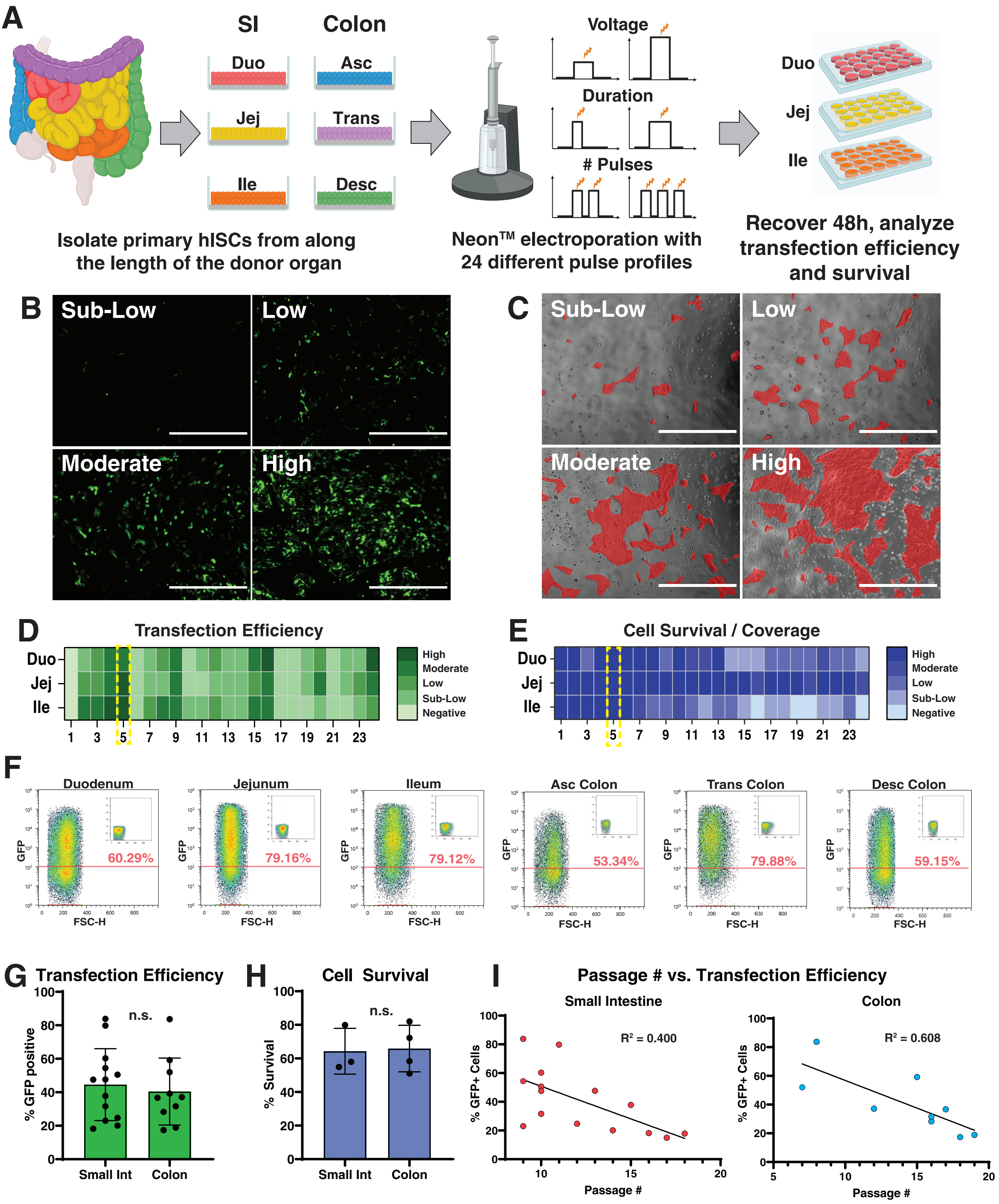

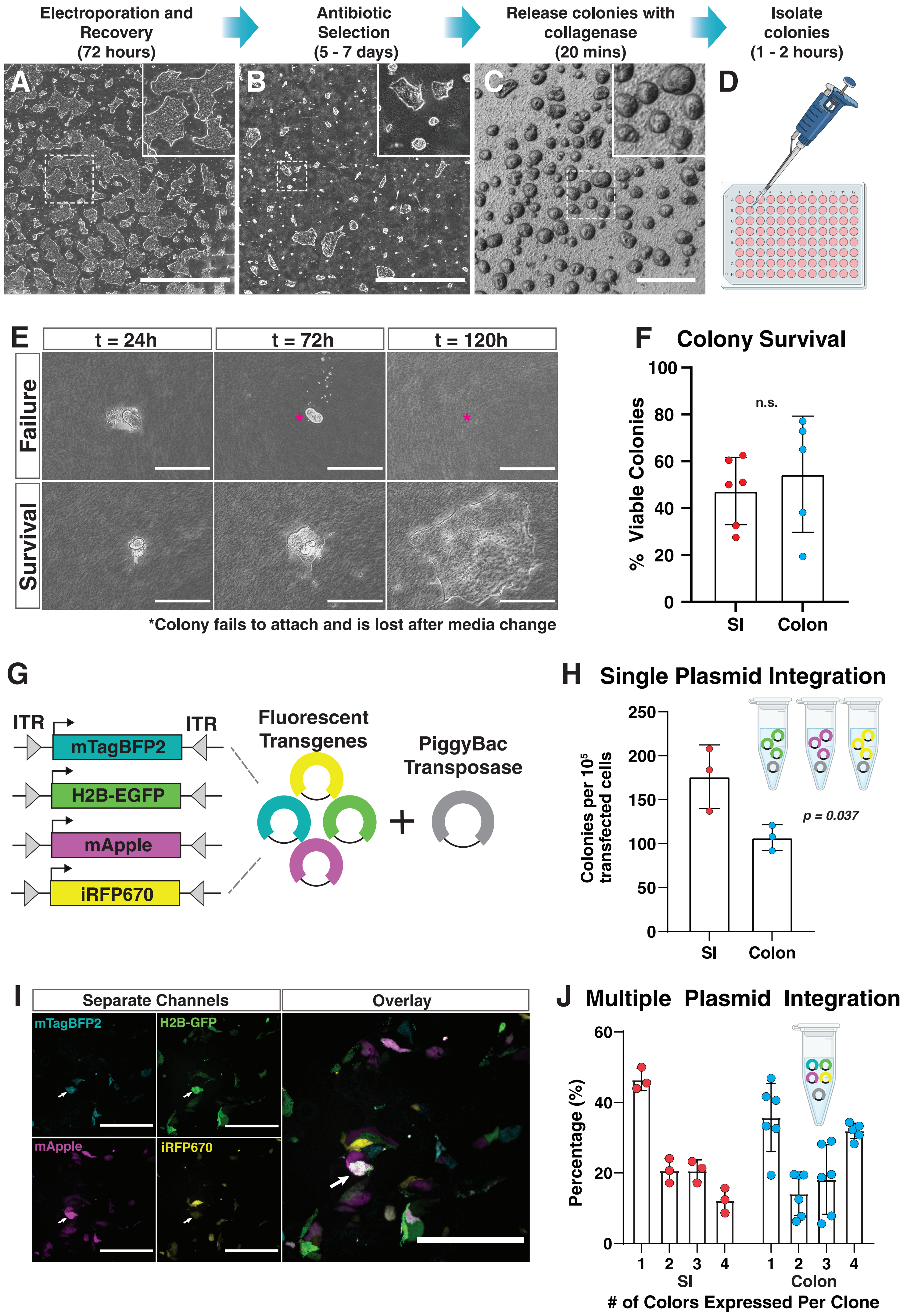

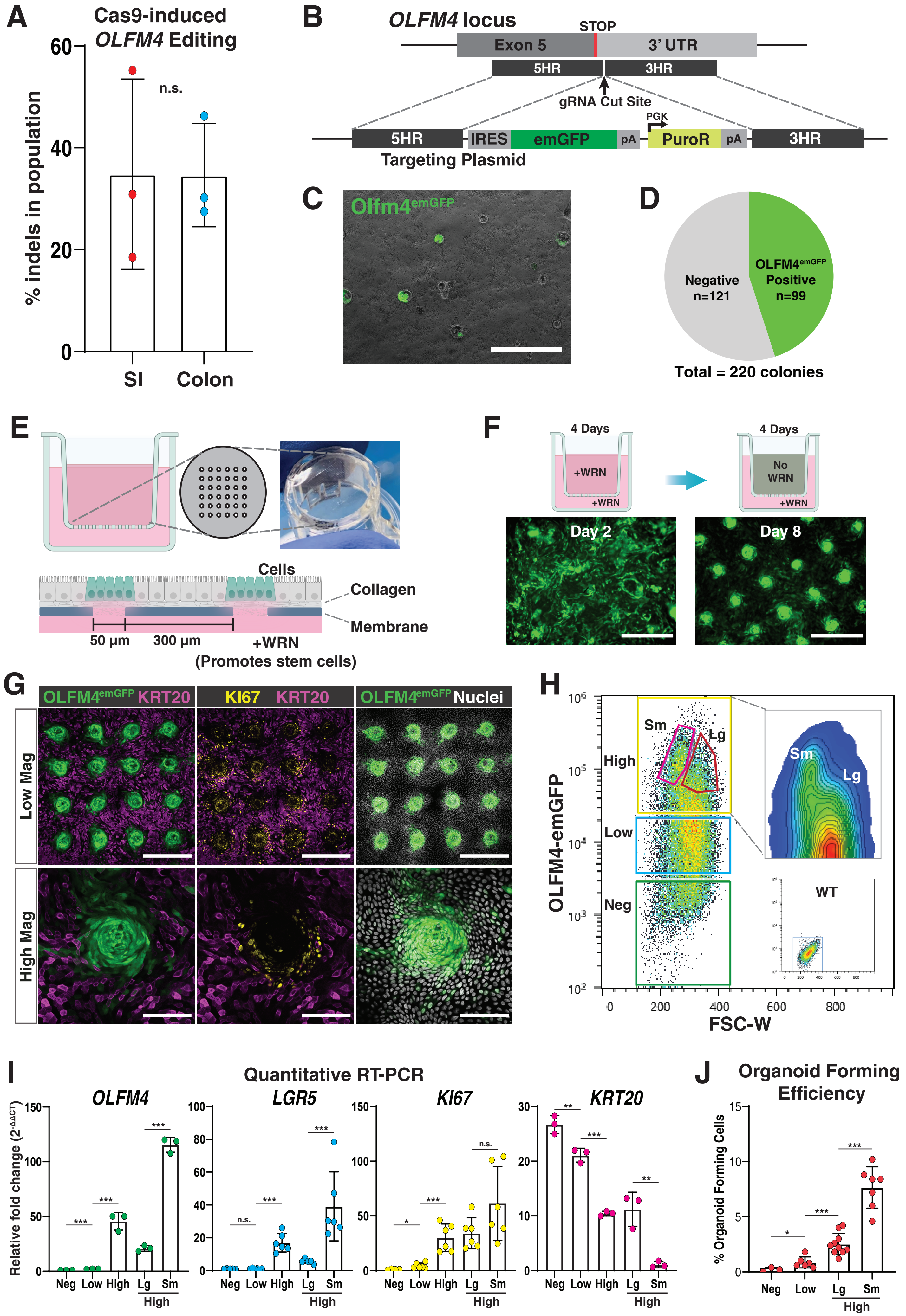
