## Supplemental Figures for "Efficient transgenesis and homology-directed gene targeting in monolayers of primary human small intestinal and colonic epithelial stem cells"

### Supplemental Table 1. Electrical pulse parameters for optimization

| Sample # | Parameters | | | Transfection Efficiency | | | Cell Viability | | |
|---|---|---|---|---|---|---|---|---|---|
| | Voltage | Length (ms) | # Pulses | Duo | Jej | Ile | Duo | Jej | Ile |
| 1 | 0 | 0 | 0 | Neg | Neg | Neg | High | High | High |
| 2 | 1400 | 20 | 1 | SubLow | Low | Moderate | High | High | High |
| 3 | 1500 | 20 | 1 | Low | Low | Moderate | Low | High | High |
| 4 | 1600 | 20 | 1 | Moderate | Moderate | High | High | High | High |
| 5 | 1700 | 20 | 1 | Moderate | High | High | High | High | High |
| 6 | 1100 | 30 | 1 | Neg | Neg | Neg | High | High | High |
| 7 | 1200 | 30 | 1 | SubLow | SubLow | Low | High | High | Moderate |
| 8 | 1300 | 30 | 1 | Low | SubLow | Moderate | High | High | Low |
| 9 | 1400 | 30 | 1 | Moderate | Moderate | Moderate | Low | High | Low |
| 10 | 1000 | 40 | 1 | Neg | Neg | SubLow | High | High | Moderate |
| 11 | 1100 | 40 | 1 | SubLow | Neg | SubLow | Moderate | High | Low |
| 12 | 1200 | 40 | 1 | Low | SubLow | Low | Moderate | High | SubLow |
| 13 | 1100 | 20 | 2 | SubLow | SubLow | SubLow | High | High | Low |
| 14 | 1200 | 20 | 2 | SubLow | SubLow | SubLow | SubLow | High | Low |
| 15 | 1300 | 20 | 2 | Moderate | Low | Moderate | SubLow | High | SubLow |
| 16 | 1400 | 20 | 2 | High | Moderate | Moderate | SubLow | Moderate | Neg |
| 17 | 850 | 30 | 2 | Neg | Neg | Neg | Low | High | SubLow |
| 18 | 950 | 30 | 2 | Neg | Neg | Neg | Low | High | SubLow |
| 19 | 1050 | 30 | 2 | SubLow | SubLow | Neg | Low | High | Neg |
| 20 | 1150 | 30 | 2 | Low | Moderate | SubLow | Low | Moderate | Neg |
| 21 | 1300 | 10 | 3 | Neg | Neg | Neg | Moderate | High | SubLow |
| 22 | 1400 | 10 | 3 | SubLow | SubLow | Neg | Low | High | SubLow |
| 23 | 1500 | 10 | 3 | SubLow | Low | SubLow | Low | High | Low |
| 24 | 1600 | 10 | 3 | High | Moderate | Low | SubLow | Moderate | Neg |

### Supplemental Table 2. CCTop off-target loci for *OLFM4* gRNA

**gRNA Sequence:** ACTTACTTAGATATCTGCAG

| Coordinates | strand | MM | target_seq | PAM | distance | | gene name | gene id |
|---|---|---|---|---|---|---|---|---|
| chr13:53050828-53050850 | + | 0 | ACTTACTT [AGATATCTGCAG] | GGG | 0 | E | OLFM4 | ENSG00000102837 |
| chr4:95984460-95984482 | - | 4 | TAACACTT [AGATATCTGCAG] | AAG | NA | - | NA | NA |
| chr6:119536897-119536919 | - | 4 | AGTAGGTT [AGATATCTGCAG] | TGG | NA | - | NA | NA |
| chr10:1497775-1497797 | + | 4 | CATTCCTG [AGATATCTGCAG] | AAG | 28840 | I | ADARB2-AS1 | ENSG00000205696 |
| chr4:158333757-158333779 | + | 4 | ATCTATGT [AGATATCTGCAG] | GGG | 18287 | I | RXFP1 | ENSG00000171509 |
| chr12:105189620-105189642 | - | 4 | GTTTAGTA [AGATATCTGCAG] | TGG | 130 | I | APPL2 | ENSG00000136044 |
| chr2:127745290-127745312 | + | 4 | GGTTAATC [AGATATCTGCAG] | CAG | 7236 | I | WDR33 | ENSG00000136709 |
| chr5:61465479-61465501 | - | 3 | ACTAATTT [GGATATCTGCAG] | TGG | 7180 | I | ZSWIM6 | ENSG00000130449 |
| chr2:219136091-219136113 | - | 4 | ACAAAATC [AGATATCTGCAG] | GGG | 10431 | I | NHEJ1 | ENSG00000187736 |
| chr3:115685944-115685966 | + | 4 | ATTTTATC [AGATATCTGCAG] | CGG | 9334 | I | GAP43 | ENSG00000172020 |
| chr21:21940614-21940636 | - | 4 | TTTTACCT [TGATATCTGCAG] | TAG | 7015 | I | AP000472.3 | ENSG00000227075 |
| chr10:114724009-114724031 | + | 4 | ATTCATTT [TGATATCTGCAG] | GAG | 33375 | - | PPIAP19 | ENSG00000228169 |
| chr18:47316063-47316085 | + | 3 | ACTTCATT [AAATATCTGCAG] | TGG | 30266 | I | CTD-2130O13.1 | ENSG00000267761 |
| chr6:81765343-81765365 | - | 4 | ATTTGCCC [AGATATCTGCAG] | CAG | 840 | - | RP5-991C6.2 | ENSG00000219702 |
| chr3:128045637-128045659 | + | 4 | ACAGATTT [TGATATCTGCAG] | AGG | 5982 | - | SEC61A1 | ENSG00000058262 |
| chr14:42710344-42710366 | - | 4 | ACAGATTT [CGATATCTGCAG] | GGG | 6689 | - | CTD-2307P3.1 | ENSG00000258394 |
| chr4:144546999-144547021 | - | 4 | ACAGATTT [TGATATCTGCAG] | AGG | 25217 | - | KRT18P51 | ENSG00000250504 |
| chr6:101110566-101110588 | + | 4 | ATTTAGAA [AGATATCTGCAG] | GAG | NA | - | NA | NA |
| chr13:39413026-39413048 | - | 4 | ATCTACTG [TGATATCTGCAG] | AAG | 34500 | I | LHFP | ENSG00000183722 |
| chr2:56417840-56417862 | - | 4 | AGAGACTT [AGTTATCTGCAG] | TAG | 31667 | - | CCDC85A | ENSG00000055813 |

### Supplemental Table 3. Primers used in this study

| Primer Name | Oligo Sequence (5' to 3') | Purpose |
|---|---|---|
| KB085_Olfm4-ScF2 | CCGTACTATGAACACCAGAACAG | PCR amplification of *OLFM4* gRNA target region for sequencing |
| KB086_Olfm4-ScR2 | CGCCATACTGAGTTTTAAGTTCC | PCR amplification of *OLFM4* gRNA target region for sequencing |
| KB026_Olfm4-SqF1 | AGCATTAACTATAACCCTTTTGAC | Sequencing primer for *OLFM4* gRNA target region |
| KB069_Olfm4-5HR-F2 | tggccgattcattaatgcagGACTGAATGCACTGAACTAAGAGG | Amplification of OLFM4 5' homology region |
| KB038_Olfm4-5HR-R | tcgaatagcttggctgcaggAGATATCTAAGTAAGTGGAGAAGAC | Amplification of OLFM4 5' homology region |
| KB039_Olfm4-3HR-F | ttaggtccctcgaggggatcGCAGGGGTGTCTAAAAGTGTG | Amplification of OLFM4 3' homology region |
| KB040_Olfm4-3HR-R | cctcttcgctattacgccagAAGGTTTCCAACTACTGCACTG | Amplification of OLFM4 3' homology region |
| KB045_emGFP-F | TCGACCTGCAGCCAAGCTATTCGAATTCCCCCCCCCCTAACGTTACTG | Amplification of emGFP coding sequence |
| KB046_emGFP-R | TGATCGGAATTGGGCTGCAGGAATTCTTACTTGTACAGCTCGTCCATGC | Amplification of emGFP coding sequence |
| KB081_IRES-SqR1 | TAACATATAGACAAACGCACACC | Screening for OLFM4-emGFP desired integration event |
| KB084_bGHpA-SqF1 | ATGCGGTGGGCTCTATGG | Screening for OLFM4-emGFP desired integration event |
| KB097_Olfm4-5LR-F1 | CTGTAGAGCTGACATTGGGTTTGC | Screening for OLFM4-emGFP desired integration event |
| KB098_Olfm4-3LR-R1 | AGGTAGCCATTCTCCCAACCTC | Screening for OLFM4-emGFP desired integration event |

### Supplemental Figure 1

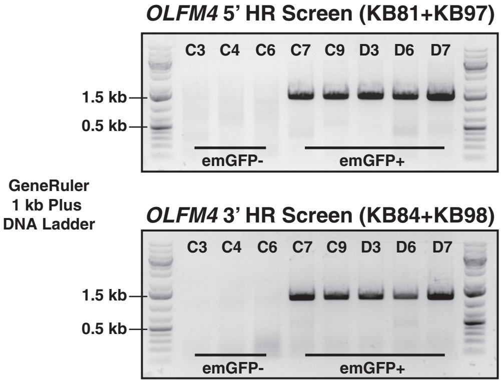


### Supplemental Figure 2

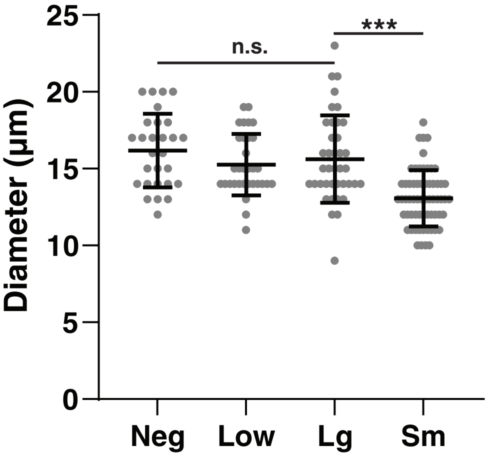
