## Supplementary material for "Efficient transgenesis and homology-directed gene targeting in monolayers of primary human small intestinal and colonic epithelial stem cells": Figure Legends

**Figure 3. An OLFM4^emGFP^ reporter generated by Cas9-mediated HDR reports proliferative stem cell populations *in vitro***. **(A)** Percentage of Cas9-induced indels at the *OLFM4* gRNA cleavage site, measured by TIDE analysis of bulk DNA collected 5 days after transfection of Cas9/gRNA complexes; *p = 0.*99, n=3 biological replicates per organ; **(B)** Schematic of the *OLFM4* insertion site and plasmid used for targeting, showing IRES-emGFP insertion in the OLFM4 3’ UTR along with a constitutively-expressed puromycin resistance gene; **(C)** Representative image of OLFM4^emGFP^ colonies following antibiotic selection; Scale bar 1000 μm; **(D)** Quantification of fluorescent OFLM4^emGFP^ colonies from Fig. 3C compared to total colony numbers; **(E)** Planar crypt-microarray (PCM) device used for hISC zonation. An array of 50 μm microholes in an impermeable membrane restricts cell access to the WRN-containing basal media reservoir, promoting stem cell maintenance only over the holes; **(F)** Overview of growth and zonation protocol for hISCs grown on PCMs. Cells are grown 4 days with WRN in both apical and basal compartments followed by 4 days with WRN in only the basal compartment; Scale bars 200 μm; (**G)** Confocal microscopy of zonated OLFM4^emGFP^ hISCs grown on PCMs, fixed and stained with antibodies for KRT20 and KI67. Nuclei counterstained with Bisbenzimide A; Scale Bars: Low Mag 400 μm, High Mag 100 μm; **(H)** FACS density plot of OLFM4^emGFP^ cells isolated from a zonated PCM, overlaid with gates used for sorting. Insets show WT negative control and a contour plot of the ‘High’ gate; **(I)** Relative gene expression measured by qRT-PCR on FACS-sorted cells obtained from each gate shown in Fig. 3H. Data normalized to ‘Neg’ for *OFLM4*, *LGR5*, and *KI67*, and to ‘High^sm^’ for *KRT20*; ** p < 0.05, ** p < 0.01, *** p < 0.001;* **(J)** Quantification of organoid formation by FACS-isolated single cells obtained from zonated PCMs; ** p<0.05, *** p<0.*001, each data point represents the percentage of cells that formed organoids following seeding of 500 single cells into a separate Matrigel™ patty.

**Supplemental Table 1.** Electrical parameters used to optimize electroporation efficiency. Twenty-four sets of parameters were tested, varying electrical voltage, pulse duration, and number of pulses. Transfection efficiency and cell viability of three biological replicates were estimated and binned into five categories (Neg, SubLow, Low, Moderate, and High). Condition #5 (1700 V, 20 ms, 1 pulse), highlighted, was used for further experiments.

**Supplemental Table 2.** CCTop output of the selected *OLFM4*-targetting gRNA, showing DNA sequences throughout the genome with homology to the gRNA sequence. First entry is the desired cleavage site in the terminal exon of *OLFM4*. Latter entries are potential off-target cleavage sites, ranked by decreasing predicted risk of cleavage, with mismatched nucleotides shown in red. gRNA was chosen such that all potential off-target sites have at least four mismatches, or three mismatches with at least one mismatch within the 12 bp “seed” region (shown in brackets), proximal to the Cas9 cleavage site.

**Supplemental Table 3.** Primers used for generating TIDE results (top), generating the *OLFM4* targeting plasmid (middle), and validating integration of the IRES-emGFP sequence (bottom).

**Supplemental Figure 1.** PCR validation of IRES-emGFP integration at the *OLFM4* locus. Three emGFP-negative clones were included for a negative control, along with five emGFP-positive clones. Primers were designed to span the homology arms used for HDR, with one primer in each screen binding to the native *OLFM4* locus outside of the homology regions and the other binding to the IRES-emGFP insert.

**Supplemental Figure 2.** Microscopy-measured cell diameters of FACS-isolated hISCs retrieved from PCM devices and embedded in Matrigel. Each data point represents a different cell. emGFP-High^Sm^ cells display significantly reduced cell size.
